# Characterization of cyanobacteria isolated from thermal muds of Balaruc-Les-Bains (France) and description of a new genus and species *Pseudo-chroococcus couteii*

**DOI:** 10.1101/2020.12.12.422513

**Authors:** C. Duval, S. Hamlaoui, B. Piquet, G. Toutirais, C. Yéprémian, A. Reinhardt, S. Duperron, B. Marie, J. Demay, C. Bernard

**Affiliations:** UMR7245 MCAM MNHN-CNRS, Muséum National d’Histoire Naturelle, CP 39, 12 rue Buffon, F-75231 Paris Cedex 05, France; Electron Microscopy Platform, Muséum National d’Histoire Naturelle, CP 39, 12 rue Buffon, F-75231 Paris Cedex 05, France; Thermes de Balaruc-Les-Bains, 1 rue du Mont Saint-Clair BP 45, 34540 Balaruc-Les-Bains

**Keywords:** cyanobacteria, thermal mud, taxonomy, polyphasic approach, morphology, ultrastructure

## Abstract

Cyanobacteria are able to synthesize a high diversity of natural compounds that account for their success in the colonization of a variety of ecological niches. Many of them have beneficial properties. The mud from the thermal baths of Balaruc-Les-Bains, one of the oldest thermal baths in France, has long been recognized as a healing treatment for arthro-rheumatic diseases. To characterize the cyanobacteria living in these muds and the metabolites they potentially produce, several strains were isolated from the water column and biofilms of the retention basin and analyzed using a polyphasic approach. Morphological, ultrastructural and molecular (16S rRNA gene and 16S-23S ITS region sequencing) methods were employed to identify nine cyanobacterial strains belonging to the orders Chroococcales, Synechococcales, Oscillatoriales and Nostocales. The combination of morphological and genetic characteristics supported the description of a new genus and species with the type species as *Pseudo-chroococcus couteii*. The high taxonomic diversity in the muds of the thermal baths of Balaruc-Les-Bains along with literature reports of the potential for bioactive metabolite synthesis of these taxa allowed us to hypothesize that some of the metabolites produced by these strains could contribute to the therapeutic properties of the muds from Thermes de Balaruc-Les-Bains.

## INTRODUCTION

Cyanobacteria belong to an ancient group of photosynthetic prokaryotes presenting a broad range of cellular strategies, physiological capacities, and adaptations that support their colonization of diverse environments worldwide. Cyanobacteria can even exist in extreme habitats and are able to settle in diverse biotopes such as hot springs. They are also known for their production of natural bioactive compounds (Demay et al. 2019), including some potent toxins (microcystins, anatoxins, saxitoxins) (Bernard et al. 2017). Some of these metabolites have been used for applications in the biotechnology and pharmaceutical fields, which has created an increased interest in the search for new isolates of cyanobacteria. Both the chemical diversity and the related bioactivity must be considered when investigating the application potential of natural products. Demay et al. (2019) concluded that among the 300 different recognized genera of cyanobacteria (referenced by the taxonomy published by Komárek et al. in 2014), 90 have been reported to produce bioactive metabolites. A few taxa are known to be prolific producers of a large set of metabolites. For example the genus *Lyngbya-Moorea*, produces 85 families of the 260 metabolites isolated so far. The majority of species have not been tested, however, and the potential for the discovery of useful natural molecules and new biosynthetic pathways from cyanobacteria remains considerable and needs to be explored.

Thermes de Balaruc-Les-Bains is one of the oldest thermal centers in France and therapeutic application of the mud, whose beneficial effects have been documented since the end of the 19th century, was recognized by the French Health System as a healing treatment for arthro-rheumatic diseases. The beneficial mud is obtained by a process of maturation in which the mud is allowed to settle naturally from a combination of hot spring water and domestic rinse water from mud treatments in a large tank. Two types of biofilms were identified within the maturation basin, one at the surface of the mud and the other on the walls of the basin. Other photosynthetic microorganisms were also described in the water column of the basin. Only two experimental studies have been carried out on mud from Thermes de Balaruc-Les-Bains. The maturation of silt from Thau Pond in Balaruc-Les-Bains was monitored for a few months in 1983, and algal development was observed ten days after formation of a mud mesocosm comprised of *Phormidium* and *Oscillatoria* Cyanobacteria, and some Diatomophyceae (Baudinat 1986). Repeated experiments in 1984 gave similar results except that algal development occurred later, after one month (Baudinat 1986). The second study was carried out in 1987 by Dupuis in experimental mesocosms for thermal mud maturation in the laboratory. In the absence of mud, there was no algal development in water from Balaruc-Les-Bains despite the addition of nutrients. In the mud mesocosms at seven weeks after the experiment started, very thick cyanobacterial mats were observed on walls and bottom, dominated by *Phormidium africanum, P. autumnale, Lyngbya* sp., *Chroococcus minor* and *Anabaena* sp. These two studies illustrated the importance of cyanobacteria as major actors in the colonization of the mud of Thermes de Balaruc-Les-Bains (Baudinat 1986 & Dupuis 1987).

In the present study, we analyzed the diversity of cyanobacteria isolated from the mud of Thermes de Balaruc-Les-Bains. Nine clonal but non-axenic strains of cyanobacteria were found and their taxonomy was verified before genomic and metabolomic analysis and bioactivities studies on their natural products were conducted. The diversity of the strains was quite high, including members of four orders, Chroococcales, Synechococcales, Oscillatoriales and Nostocales.

## MATERIALS AND METHODS

### Sampling

Samples for strain isolation were collected twice a month from April 28 to October 13, 2014 (Table 1) from the thermal basins of Balaruc-Les-Bains (43°26’44.0”N 3°40’29.6”E).

**Table 1.** List of cyanobacteria strains isolated from the retention basin of Thermes of Balaruc-Les-Bains and their corresponding strain numbers, habitat (water column or biofilms) and sampling dates. PMC: Paris Museum Collection.

| Order | Species | Strain number | Habitat | Sampling date |
| --- | --- | --- | --- | --- |
| Chroococcales | *Pseudo-chroococcus couteii* | PMC 885.14 | Epilithic biofilm | 23/06/2014 |
| Synechococcales | *Leptolyngbya boryana* | PMC 883.14 | Mud surface | 12/05/2014 |
| Oscillatoriales | *Planktothricoides raciborskii* | PMC 877.14 | Mud surface | 26/05/2014 |
|  | *Laspinema* sp. | PMC 878.14 | Mud surface | 23/06/2014 |
|  | *Microcoleus vaginatus* | PMC 879.14 | Epilithic biofilm | 26/05/2014 |
|  | *Lyngbya martensiana* | PMC 880.14 | Epilithic biofilm | 23/06/2014 |
| Nostocales | *Nostoc* sp. | PMC 881.14 | Epilithic biofilm | 23/06/2014 |
|  | *Aliinostoc* sp. | PMC 882.14 | Epilithic biofilm | 23/06/2014 |
|  | *Calothrix* sp. | PMC 884.14 | Mud surface | 12/05/2014 |

The 500 m^3^ maturation basin (Figure 1a, c and d) received the muddy water from the rinsing of the patients. The slurry was a mixture of water from the thermal springs along with domestic hot water containing the mud particles, which settle to the bottom (Figure 1b and c).

**Figure 1.**
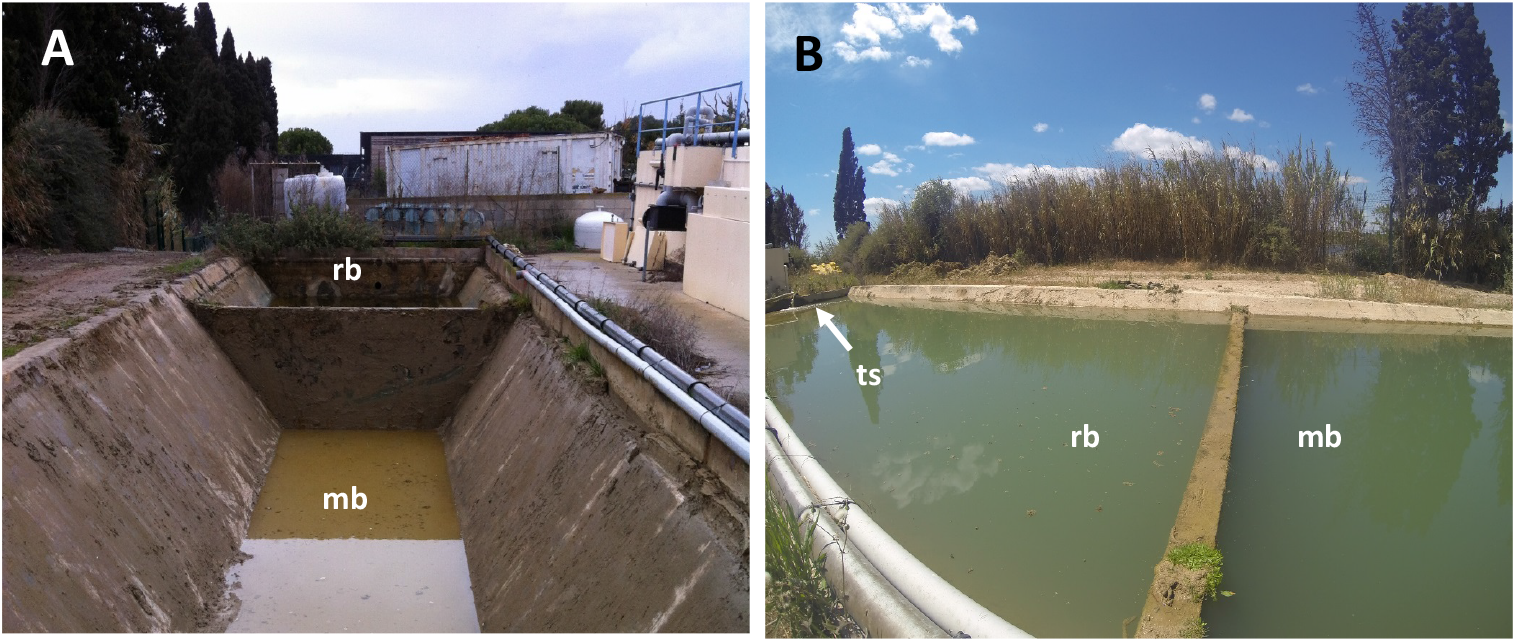
Retention basin of the Thermes de Balaruc-Les-Bains. Views of the mud maturation basin (mb), the water retention basin (rb) and the thermal spring water supply (ts) on February 28t^h^, 2014 (A) and on May 12^th^, 2014 (B).

The water in the retention basin was stirred by the continuous influx of rinse water, and the level was kept above the mud by draining excess water to the outside to prevent overflow (Figure 1d). The proper temperature and light levels and nutrient inputs were maintained to ensure that the mud maturation process was optimal. Three types of sampling were carried out from the maturation basin: (*i*) from the biofilm covering the mud, (*ii*) from the epilithic biofilm on the walls of the basin where algae development was more extensive and (*iii*) from the water surface using a 20-μm membrane filter.

### Cyanobacterial culture conditions and strain isolation

Samples were inoculated on solid medium (5 or 10 g·L^−1^ agar) Z8 and Z8-salts (Rippka 1988). Isolations were carried out by repeated transfers (at least three times) of single cells or filaments on solid or liquid media under an inverted microscope (Nikon ECLIPSE TS100). Viable clones were then cultured in 25 cm^2^ culture flasks (Nunc, Roskilde, Denmark) containing 10 mL of Z8. Cyanobacterial strains were maintained in the Paris Museum Collection (PMC) at 25°C, using daylight fluorescent tubes providing an irradiance of 12 μmol photons·m^−2^·s^−1^, with a photoperiod of 16h light/8h dark. Isolated strains and cultures were all monoclonal and non-axenic.

### Morphological analyses

Morphological analyses of cyanobacterial strains were carried out using an Axio ImagerM2 Zeiss microscope. Photographs of the specimens were taken with an AxioCam MRc digital camera coupled to the microscope and images were processed using the ZEN software (Zeiss). The cell and filament width and length, morphology, color and motility were determined. Strains were identified morphologically using the updated taxonomic literature (Komárek and Anagnostidis 2005; Komárek and Anagnostidis 2007; Komárek and Johansen 2015).

### Pigment analysis

Pigment analysis was performed on each strain in triplicate. Ten mL aliquots of culture in exponential growth were filtered through Whatman GF/F filters (Ø 0.7 µm) and freeze-dried. The lipophilic carotenoids and chlorophylls were extracted, analyzed and quantified by HPLC-DAD as described (Ras et al., 2008). The water soluble phycobiliproteins were analyzed following a Standard Operating Procedure (Yéprémian et al. 2017).

### Ultrastructural analyses

Cyanobacterial strains were imaged by scanning electron microscopy (SEM) and transmission electron microscopy (TEM) as described by Parveen et al. (2013), with modifications. Cells or filaments harvested by centrifugation from a growing culture were fixed with 2% (v/v) glutaraldehyde, 2% (v/v) formaldehyde, 0.18 M sucrose, and 0.1% picric acid in 0.1 M Sorensen phosphate buffer (SPB; pH 7.4) for 1 h, at room temperature. The specimens were then washed three times with SPB and post-fixed with 1% osmium tetroxide for 1 h. Afterwards, they were washed with distilled water before being dehydrated in a microscopy-grade ethanol series (30%, 50%, 70%, 90% and 100%), with agitation and centrifugation. For SEM, the samples were pipetted onto glass coverslips on SEM stubs and air dried. They were coated with platinum (Leica EM ACE600 coater) and examined using a scanning electron microscope (Hitachi SU3500, Japan). For TEM, the samples were embedded in epon resin and sectioned at 0.5 μm using an ultra-microtome (Reichert-Jung Ultracut) with a diamond knife and transferred onto 150-mesh copper grids. The prepared sample grids were stained with 2% uranyl acetate in 50% ethanol for 15 min and washed three times in 50% ethanol and twice in distilled water. The copper grids were then dried, examined with a transmission electron microscope (Hitachi HT-7700, Japan), and photographed with a digital camera (Hamamatsu, Japan).

### Molecular and phylogenetic analyses

DNA was extracted from cyanobacterial strains using the ZymoBIOMICS DNA mini kit (Zymo Research, CA) following the manufacturer’s protocol. Cells were lysed using a Qiagen bead mill for 6 min at maximum speed (30 Hz, 1800 oscillations per minute). The amplification of the 16S rRNA-encoding gene and the 16S-23S internal transcribed spacer (ITS) region was done with the primers and PCR programs described in Cellamare et al. (2018) and Gama et al. (2019), respectively. All PCR reactions were carried out using a GeneTouch thermocycler (Bioer Technology, Hangzhou, China). The extracted and purified PCR products were then sequenced by Genoscreen (Lille, France). Sequences were assembled and corrected using the MEGA version 7.0 software (Kumar et al. 2016) and aligned using MAFFT online service (Katoh, et al. 2019). For the 16S rRNA gene, partial sequences with a minimum of 1398 base pairs (bp) from the complete gene sequence (1550 bp) were obtained. For the 16S-23S ITS, partial sequences were obtained with a minimum of 174 bp. Phylogenetic analyses were performed according to three methods: maximum parsimony (MP), neighbor joining (NJ), and maximum likelihood (ML) using the MEGA version 7 software with an equal-to-1000 iterations.

For 16S rRNA gene phylogeny, an overall alignment (n = 129 sequences) was generated, including the newly produced sequences and reference sequences available in GenBank representing Oscillatoriales, Synechococcales, Nostocales and Chroococcales. The selected sequences were all longer than 1200 bp. *Gloeobacter violaceus* PCC 7421 was chosen as outgroup. A cut-off value of 95% of 16S rRNA gene sequence identity was used for genus definition (Komárek 2010).

For the *Chroococcus*-like strain, the 16S-23S ITS phylogeny was performed with representative strains from *Chroococcus*-like, *Inacoccus, Cryptococcum, Gloeocapsa* (*syn. Limnococcus*) and *Chroococcus*, whose sequences are available in Genbank and (Gama et al. 2019). An overall alignment (n = 21) was generated with the selected sequences, all longer than 1494 bp. *Microcystis aeruginosa* PMC 728.11 was chosen as an outgroup. The generated 16S-23S ITS sequences were used for determination of secondary structure. The conserved regions (D1–D1’) were analyzed using the Mfold WebServer (Zuker 2003). Default settings were used, except for the structure drawing mode where the natural angle was selected. Transfer RNA genes were found using tRNAscan-SE 2.0 (Lowe and Chan 2016). The nucleotide sequences reported in this study have been deposited in the National Center for Biotechnology Information database. GenBank accession numbers are listed in the Supplementary table (Table S1).

## RESULTS AND DISCUSSION

Nine clonal strains of cyanobacteria, living in the mud maturation basin of the Thermes de Balaruc-Les-Bains, were isolated (Table 1). These organisms were representative of the algal communities in (Baudinat 1986; Dupuis 1987). Dupuis (1987) showed that *Phormidium* was the most abundant genus in muds of Balaruc-Les-Bains with the species *Phormidium africanum* as dominant and *P. autumnale, Lyngbya sp*. and *Chroococcus* as accompanying species. Baudinat (1986), also showed that the cyanobacteria were dominant, mainly represented by *Phormidium* and *Oscillatoria*. A preliminary study done in 2014 during the mud maturation period showed that the cyanobacterium, *Planktothricoides raciborskii*, was also quite dominant (unpublished data). Other taxa such as *Laspinema* sp., *Leptolyngbya boryana* and *Nostoc* sp. were observed. The dominance of *Laspinema* sp. was observed on one sampling date, corresponding to a slight increase in salinity of the thermal water. Other cyanobacterial taxa were observed such as *Calothrix* sp. in the mud and *Aliinostoc* sp., *Chroococcus*-like and *Microcoleus vaginatus* on the walls of the maturation basin. As *Planktothricoides raciborskii* is morphologically very similar to *Phormidium*, we concluded that the dominance of the genus *Phormidium* has been stable since the 1980s.

To characterize the nine cyanobacterial isolates, we used morphological, ultrastructural and 16S rRNA and 16S-23S internal transcribed spacer (ITS) gene sequence analyses. Thus, eight cyanobacteria were firmly identified at the genus or species level, while one remained unidentified (PMC 885.14). The phylogenetic trees based on the 16S rRNA gene sequences shared well-supported nodes at the order level regardless of whether the method used was MP, NJ or ML (see the ML tree, Fig. 2, and the three consensus phylogenies, Fig. S2). Some isolates were clearly identified at the species level, while others were only assigned to the genus level because of the large species diversity within the given genus.

**Figure 2.**
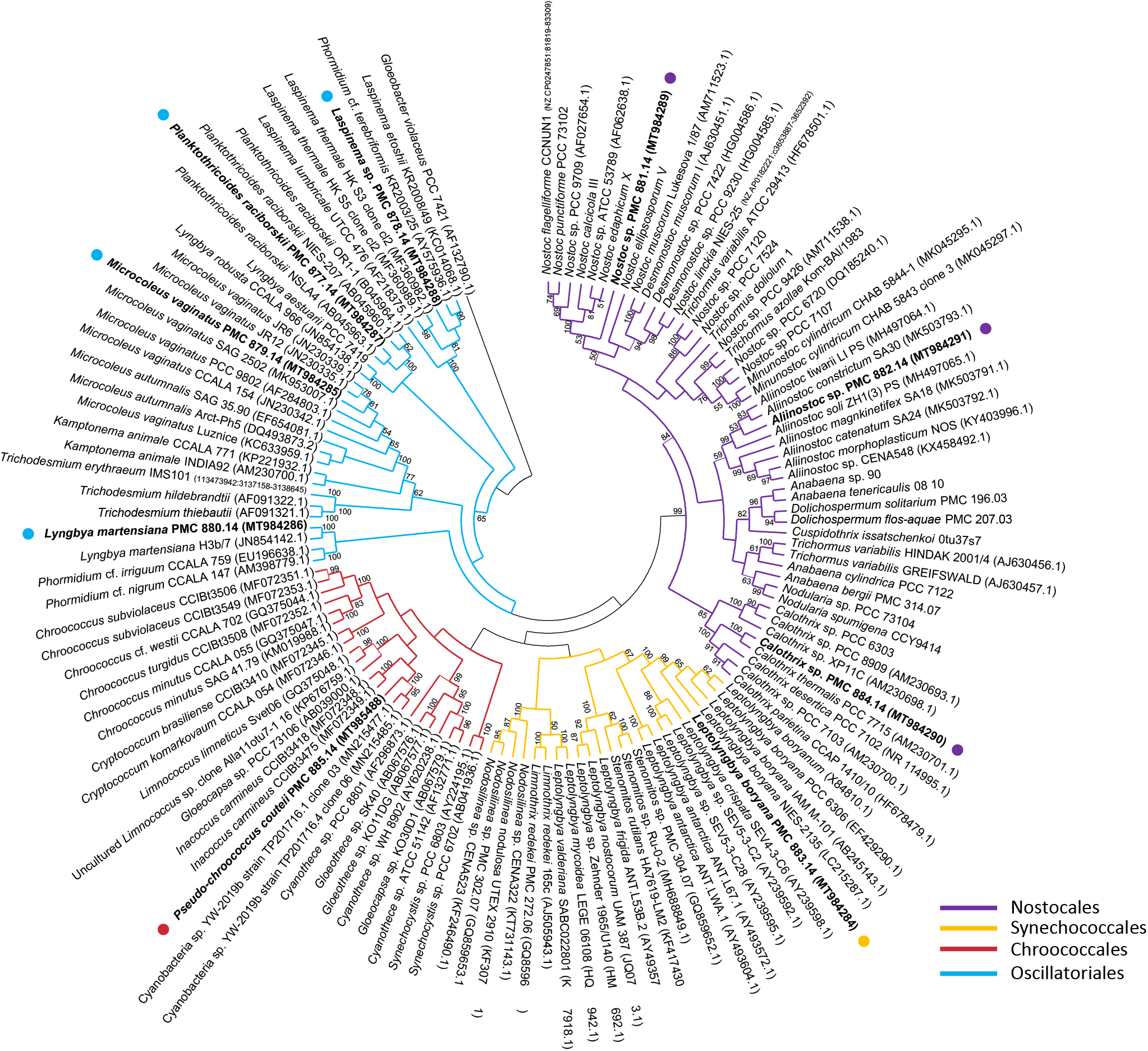
Maximum likelihood phylogenetic tree based on 16S rRNA gene sequences of representative cyanobacteria strains (129 sequences, 1446 aligned nucleotide positions, GTR+G+I model) belonging to the orders Oscillatoriales, Nostocales, Chroococcales, Synechococcales and the studied strains form Thermes de Balaruc-Les-Bains (in bold). *Gloeobacter violaceus* was used as outgroup. Numbers above branches indicate bootstrap support (>50% shown) based on 1000 replicates.

The nine new strains belonged to the orders Chroococcales: *Pseudo-*c*hroococcus couteii* (PMC 885.14); Synechococcales: *Leptolyngbya boryana* (PMC 883.14); Oscillatoriales: *Planktothricoides raciborskii* (PMC 877.14), *Laspinema* sp. (PMC 878.14), *Microcoleus vaginatus* (PMC 879.14), *Lyngbya martensiana* (PMC 880.14) and Nostocales: *Nostoc* sp. (PMC 881.14), *Aliinostoc* sp. (PMC 882.14), *Calothrix* sp (PMC 884.14). Identification numbers and habitats of the strains are given in Table 1.

### Characterization of strains belonging to the Chroococcales

Among the nine studied strains, one unidentified strain displayed a *Chroococcus*-like morphology (PMC 885.14) and belonged to the Chroococcales; but, the 16S rRNA sequence diverged from known *Chroococcus* sequences by more than 5%, supporting the idea that it was a new genus.

***Pseudo-chroococcus couteii*** (C. Duval, S. Hamlaoui, C. Bernard gen. nov., sp. nov., 2020) (PMC 885.14) *Pseudo-chroococcus couteii* (PMC 885.14) was isolated from an epilithic biofilm in the mud maturation basin of Thermes de Balaruc-Les-Bains (Table 1). Examination of samples in the field showed single cells or small colonies of 2-8 cells surrounded by dense, colorless and slightly lamellate mucilaginous envelopes (Fig. 3). Under both natural and laboratory conditions, cells were spherical or hemispherical 14-18 µm in diameter with pale-green, blue-green or grey homogeneous to granulate cell contents (Fig. 3). Cell division occurred by binary fission in three or more planes or irregularly in old colonies. These morphological features are typical of the genus *Chroococcus*. TEM ultrastructure imaging showed a dense mucilaginous layer surrounding four grouped cells (Fig. 3). Thylakoids were fasciculate, as found in *Chroococcus* species.

**Figure 3.**
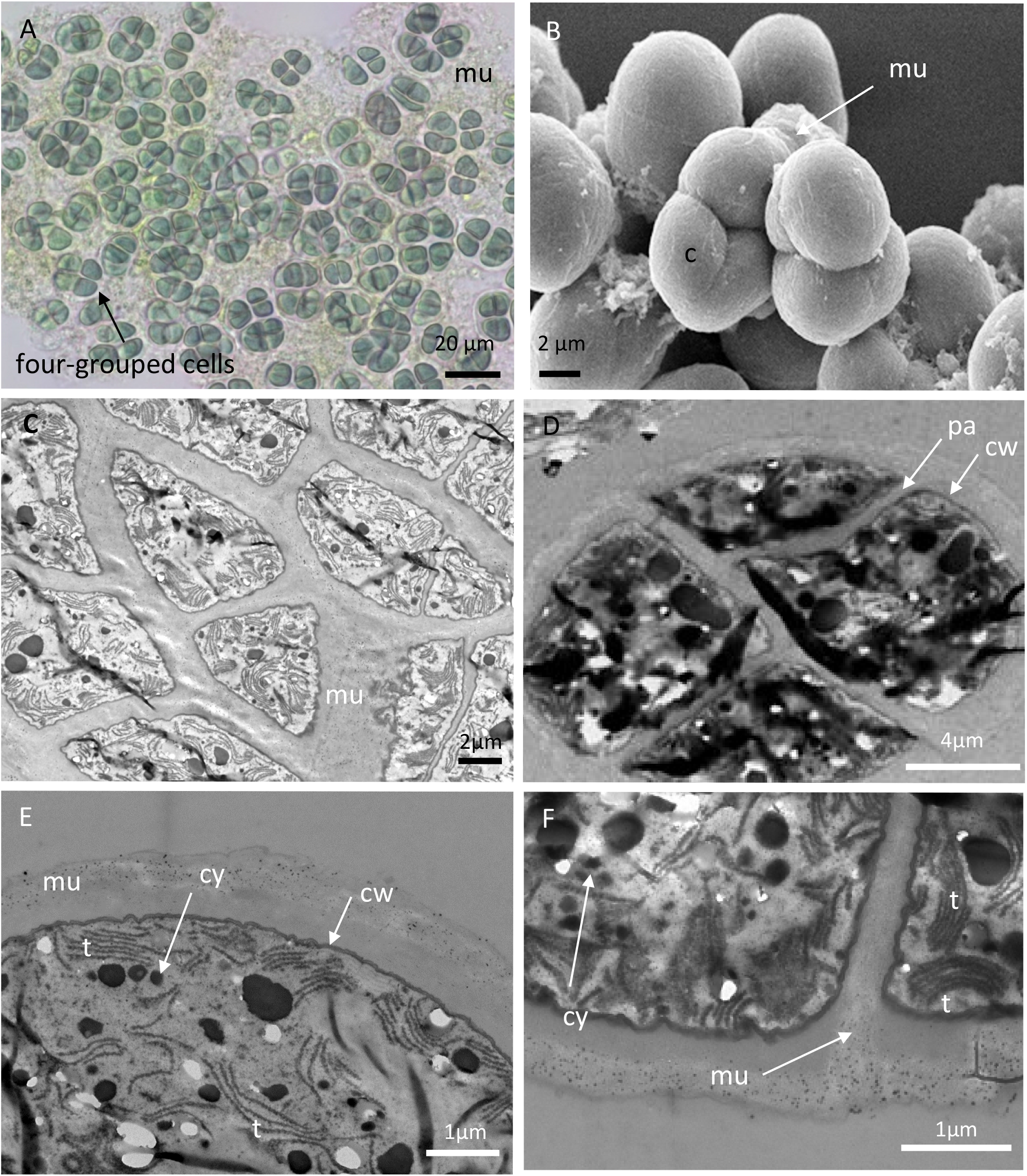
Light microscopy (A), SEM (B) and TEM (C-F) micrographs of *Pseudo-chroococcus couteii* (PMC 885.14) from Thermes de Balaruc-Les-Bains. A: Light microscopy of a packet-form colony, four-celled groups of spherical, very small cells, brownish color with dense mucilaginous sheath around cells. B: external observation of a packet-form colonies C, D, E, F.: the four-grouped cells were surrounded by a dense mucilage. Cells details showing a fasciculate arrangement of thylakoids. Abbreviations: c: colony of four-grouped cells, cy: cyanophycine granules, cw: cell wall, pa: cells partition, mu: mucilage, t: thylakoids.

The 16S rRNA gene sequence analyses showed that the *Pseudo-chroococcus couteii* strain PMC 885.14 had 93% and 92% sequence identity with *Inacoccus* and *Cryptococcum* gen. nov. strains respectively (data not shown). The 16S rRNA sequence data also revealed the polyphyletic nature of the genus *Chroococcus*. The ML phylogenetic tree of the 16S rRNA and the 16S-23S ITS gene sequences (Fig. 4 and Fig. 5) of *Chroococcus* strains produced three main clusters, *Chroococcus*-like, *Inacoccus*/*Cryptococcum* and *Limnococcus*/*Gleocapsa*, separated from the *Chroococcus sensu stricto* cluster. The first cluster was mainly composed of three strains of *Inacoccus carmineus* from concrete of Santa Virgínia Park in Brazil and two strains of *Cryptococcum* from thermal springs and soil. The second cluster grouped *Limnococcus limneticus* and *Gleocapsa* sp. 16S rRNA gene sequences. The third *Chroococcus*-like cluster consisted of two sub-clusters: one composed of three marine strains isolated from surface sediments in New Zealand and Hawaii, and one composed of PMC 885.14 (this study) and four freshwater strains from China. The two sub-clusters were well-supported (100% bootstrap support).

**Figure 4.**
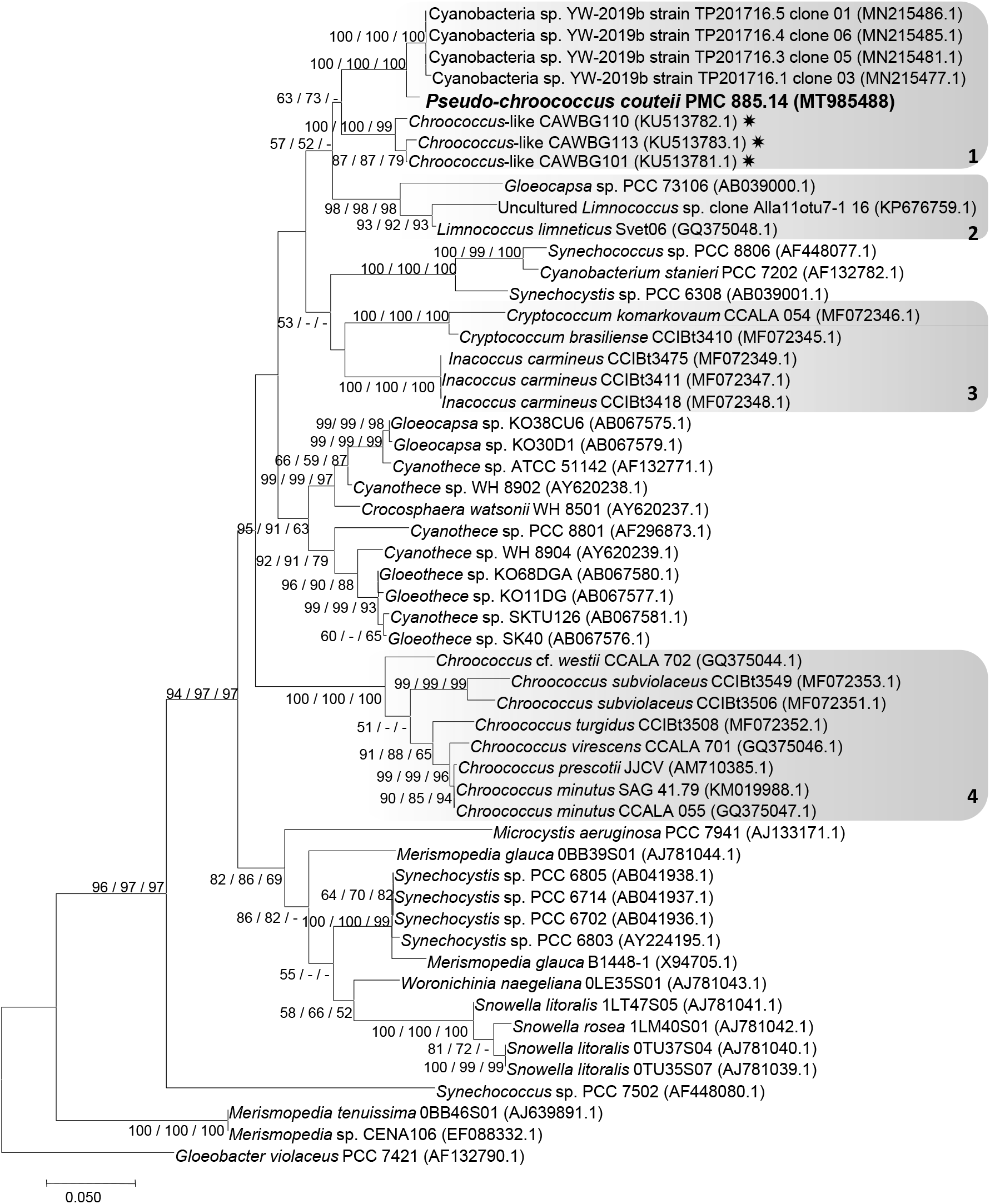
Consensus phylogenetic tree (Maximum likelihood tree presented) based on 16S rRNA-encoding gene sequences (54 sequences, 705 aligned nucleotide positions, GTR+G+I model) of representative cyanobacteria strains belonging to the order Chroococcales. Thermes de Balaruc-Les-Bains’s strain is indicated in bold, other species and sequences were obtained from Genbank. *Gloeobacter violaceus* PCC 7421 was used as an out-group. Numbers above branches indicate bootstrap support (>50%) from 1,000 replicates. Bootstrap values are given in the following order: maximum likelihood / neighbor joining / maximum parsimony. (✷ marine strains; Cluster 1 : *Chroococcus*-like, 2 : *Limnococcus*/*Gloeocapsa*, 3 : *Inacoccus*/*Cryptococcum*, 4 : *Chroococcus sensu stricto*)

**Figure 5.**
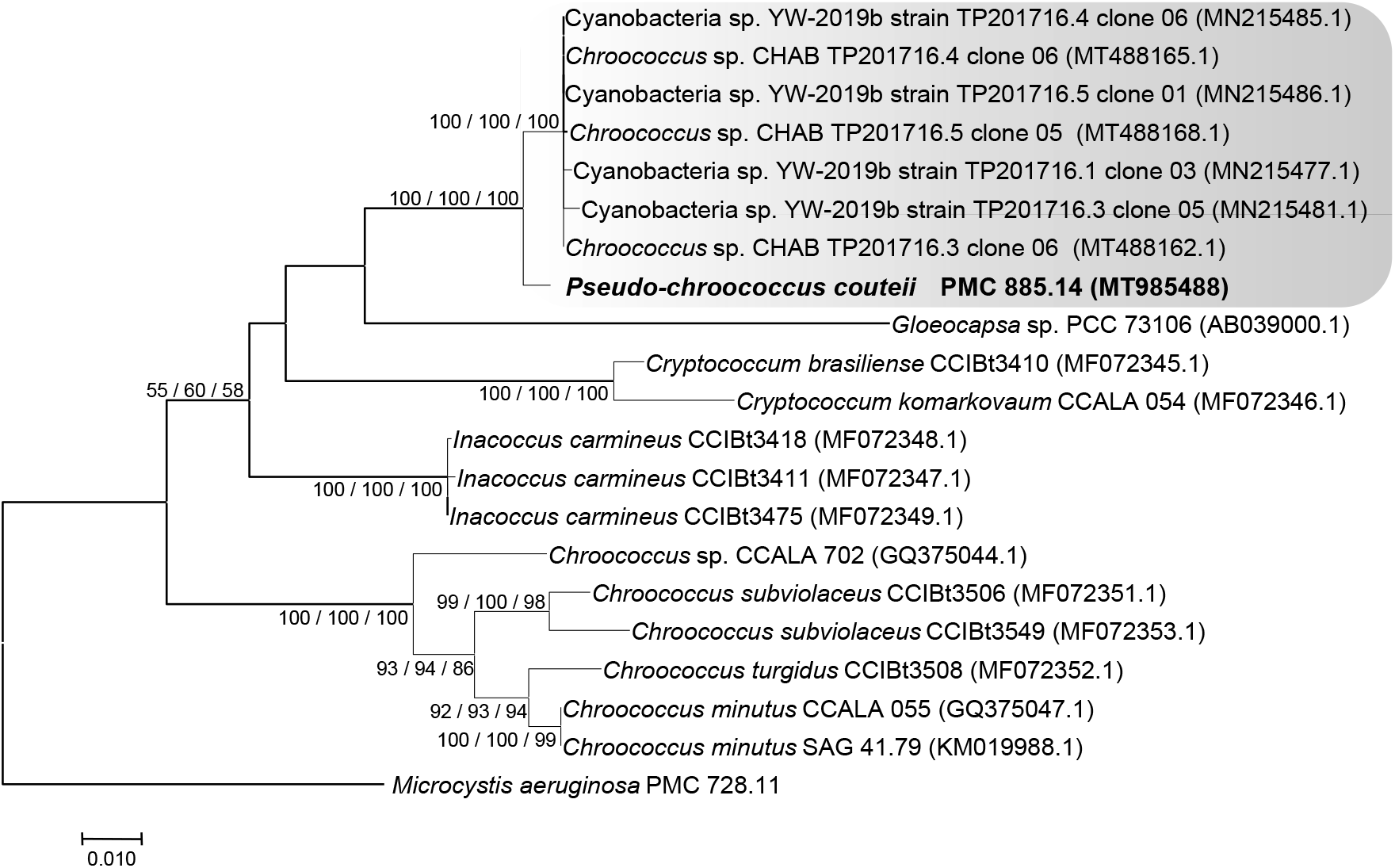
Consensus phylogenetic tree (Maximum likelihood tree presented) based on 16S-23S ITS gene sequences (21 sequences, 1915 aligned nucleotide positions, GTR+G+I model) of representative cyanobacteria strains belonging to the orders Chroococcales, Thermes de Balaruc-Les-Bains’s strains (in bold), other species were obtained from Genbank. *Microcystis aeruginosa* PMC 728.11 was used as an outgroup. Numbers above branches indicate bootstrap support (>50%) from 1000 replicates. Bootstrap values are given in the following order: maximum likelihood / neighbor joining / maximum parsimony.

Substantial differences were observed among the secondary ITS structures (D1-D1’ helices) of the three genera previously mentioned (Fig. 6). The pattern and the number of nucleotides (nt) in the D1-D1’ region of *Chroococcus sensu stricto* (58 nt), *Inacoccus carmineus* (63-64 nt), *Cryptococcum* (56-59 nt) and *Chroococcus*-like (48-49 nt) were different. The pattern of the terminal portion of *Pseudo-chroococcus couteii* strain (PMC 885.14) from Thermes de Balaruc-les-Bains was different from that of freshwater strains from China (YW-2019b strain TP20176.5 clone 0.1 and CHA TP20176.4 clone 0.6), with a single loop of 18 nt for *Pseudo-chroococcus couteii* and two loops of 4 nt and 9 nt for the Chinese strains (Fig. 6). A unilateral bulge of 7 nt (CAAUCC) was present in the three strains of the *Chroococcus*-like cluster.

**Figure 6.**
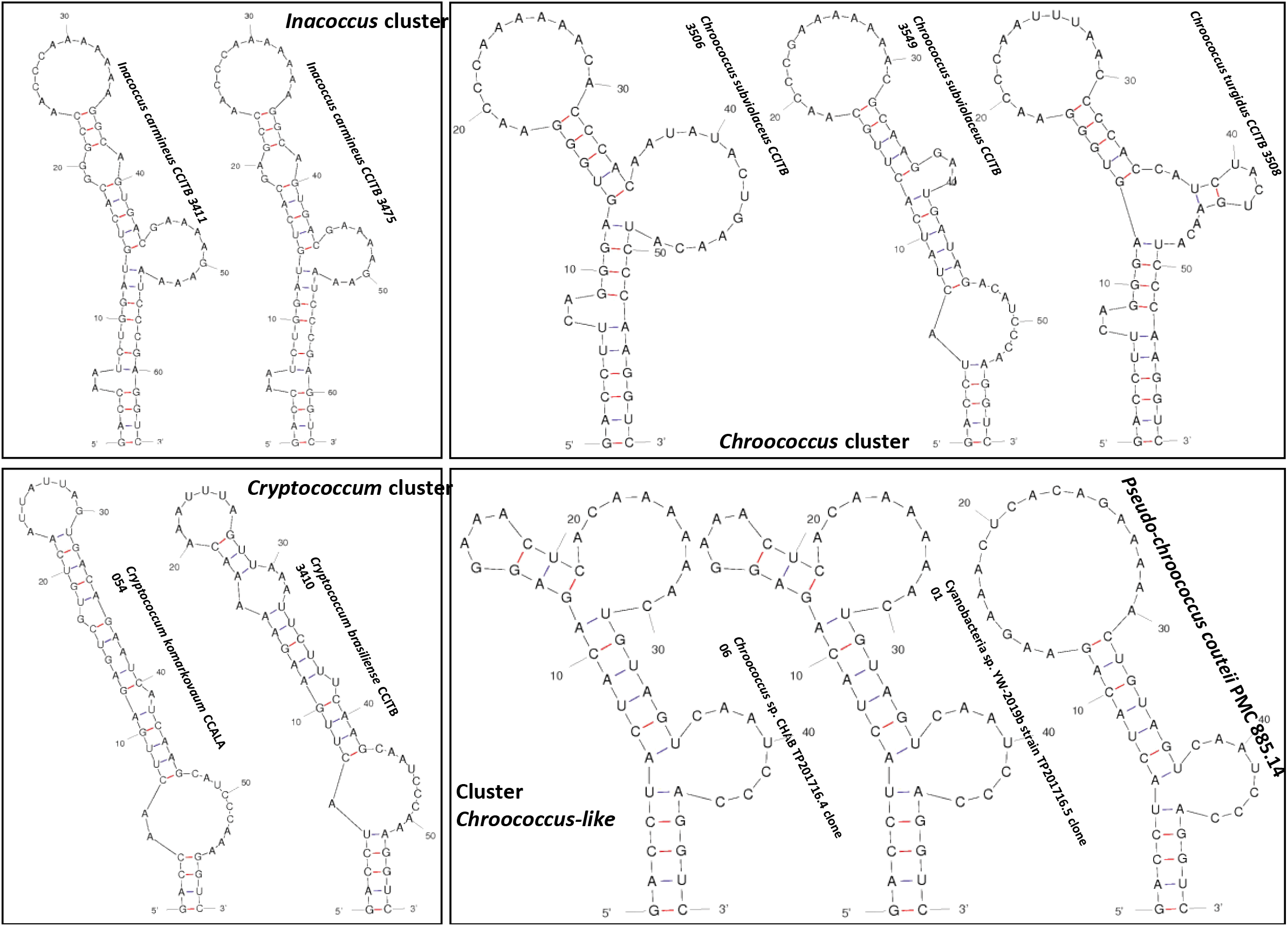
16S-23S ITS secondary structure of the D1-D1’ helices of *Inacoccus, Cryptococcum, Chroococcus* and *Chroococcus*-like (syn. *Pseudo-chroococcus*) cluster.

To the best of our knowledge, only four studies to date have examined phylogenetic relationships (16S rRNA) within the genus *Chroococcus* (Komărkovă et al. 2010; Roldán et al. 2013; Wood et al. 2017; Gamma et al. 2019). The strains studied were all morphologically similar to *Chroococcus* species in cell shape and organization (Table S2). However, they differed by the color of the sheaths or mucilage (*e*.*g*., the mucilage of *Chroococcus* was usually hyaline or yellowish whereas that of *Limnococcus* was described as red or brown) (Gama et al. 2019; Table S2). Wood et al. (2017) stated that within the 51 *Chroococcus*-like 16S rRNA sequences, only five were obtained from isolates, limiting phylogenetic investigations.

However, this study included the new 16S rRNA sequences available in GenBank and clearly demonstrated that the *Chroococcus*-like cluster was well-supported and should be described as a new genus.

***Pseudo-chroococcus couteii*** by C. Duval, S. Hamlaoui, C. Bernard gen. nov., sp. nov.

### Diagnosis

Solitary cells or small colonies of 2, 4 or 8 cells surrounded by dense, colorless and slightly lamellate mucilaginous envelopes. Cells, spherical or hemispherical 14-18 µm in diameter with pale-green, blue-green or grey homogeneous to granulate cell contents. Cell division occurring by binary fission in three or more planes, or irregularly in old colonies.

### Holotype

A formaldehyde-fixed, cryopreserved sample of the strain PMC 885.14 was deposited at the Paris Museum Collection (PMC), Paris, France.

### Type strain

A live culture was deposited at the Paris Museum Collection (PMC) as PMC 885.14, with GenBank accession no. MT985488 for the 16S-23S rRNA gene sequence.

### Type locality

epilithic biofilm, retention basin of the thermal springs at Balaruc-Les-Bains, France (GPS: 43°26’44.0”N 3°40’29.6”E).

### Etymology

the name of the genus, *Pseudo-chroococcus*, refers to the cyanobacterium morphology (colonial) like *Chroococcus sensu stricto*, but, clearly phylogenetically separate in two different clades; *couteii* = named in honor of Prof. Alain Couté, a French phycologist at the National Museum of Natural History in France.

### Characterization of strains belonging to the Synechococcales

Among the nine studied strains, one strain *Leptolyngbya boryana* (PMC 883.14) belonged to the Synechococcales.

#### *Leptolyngbya boryana* (Anagnostidis et Komárek, 1988)

*Leptolyngbya boryana* (PMC 883.14) was isolated from mats covering the surface of the basin of Thermes de Balaruc-Les-Bains (Fig. 1, Table 1). Examination of samples in the field showed a pale-green, diffluent thallus forming mats (Fig. 7). Under laboratory conditions, the filaments were straight to slightly curved and densely entangled with thin, firm, colorless sheaths. Trichomes were pale to bright blue-green, cylindrical, unbranched, thin, < 3 µm wide and strongly constricted at the ungranulated cross-walls (Fig. 7), motile (hormogonia) or immotile, or with indistinct trembling. Cells were moniliform, shorter than wide, 1.5-2.4 μm wide and 0.9-2.0 µm long with a length:width ratio of 0.76 and well-separated from one another by cross-walls. Cell contents were pale blue-green and homogeneous without gas vacuoles. Apical cells were rounded without calyptra (Fig. 7). A sheath with a thin transparent zone covered the trichome and was clearly visible by TEM. The thylakoids were parietally arranged, concentric and parallel to the cell wall (Fig. 7).

**Figure 7.**
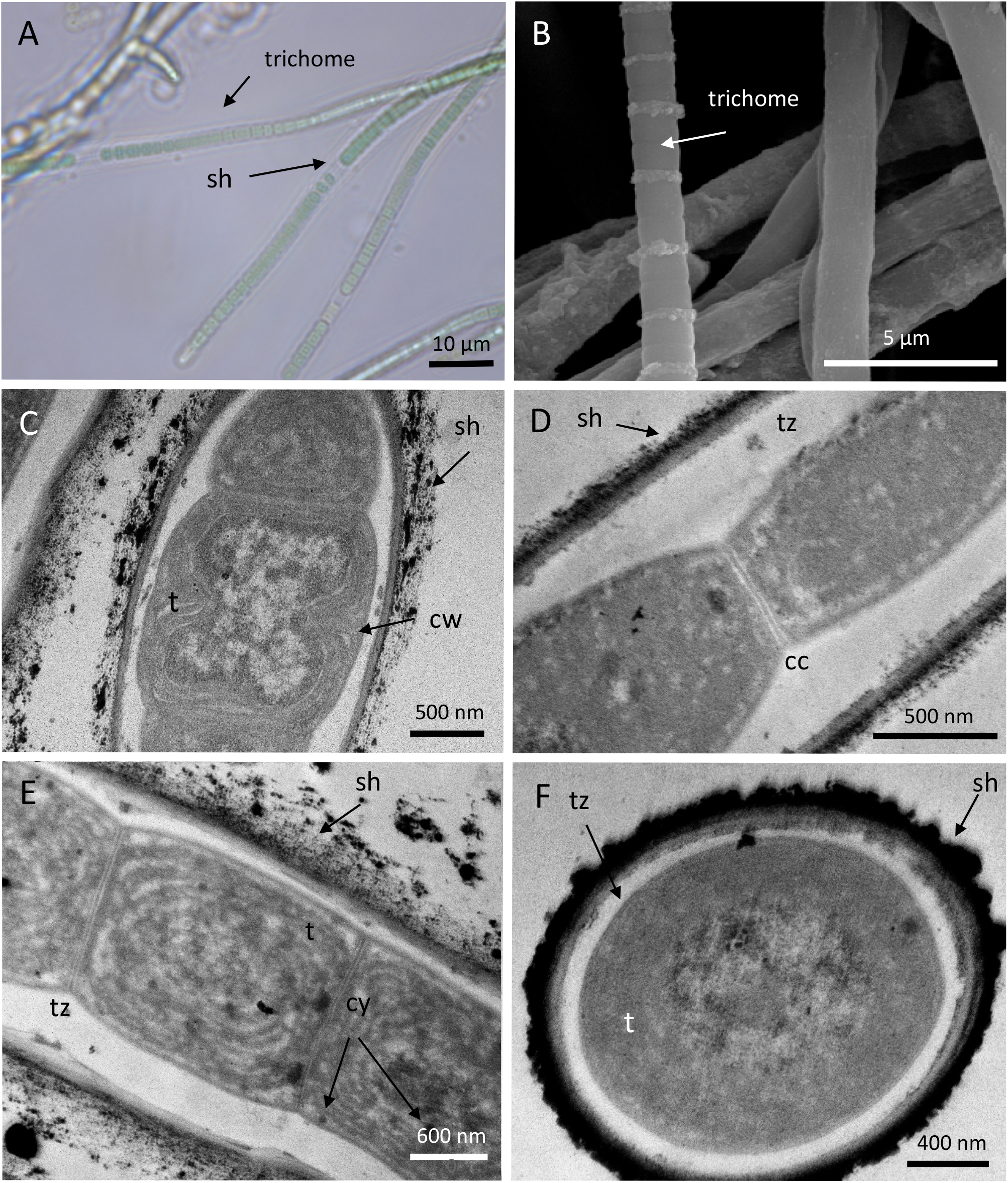
Light microscopy (A, B), MEB (C) and TEM (D-F) micrographs of *Leptolyngbya boryana* (PMC 883.14) from Thermes de Balaruc-Les-Bains. A: entangled filaments and trichomes with sheath. B: observations of trichomes with probable exudates localized on cell junctions. Longitudinal (C-E) and cross sections (F) of trichomes with sheaths showing the ultrastructural details. Abbreviations: cc: cross-wall constriction, cm: cytoplasmic membrane, cy: cyanophycin granules, t: thylakoids, tz: transparent zone.

Results based on 16S rRNA gene sequence homology (BLAST), showed that the *Leptolyngbya* strain (PMC 883.14) shared 100% sequence identity with other *Leptolyngbya* strains (data not shown). This result was confirmed by the phylogenetic tree (Fig. 8) where the strain PMC 883.14 was grouped in a well-supported clade including the *Leptolyngbya boryana*, cluster with one *Leptolyngbya foveolarum* sequence (X84808.1) and one *Leptolyngbya boryanum* sequence (X84810.1). The cluster was mainly comprised of strains isolated from benthic freshwater environments.

**Figure 8.**
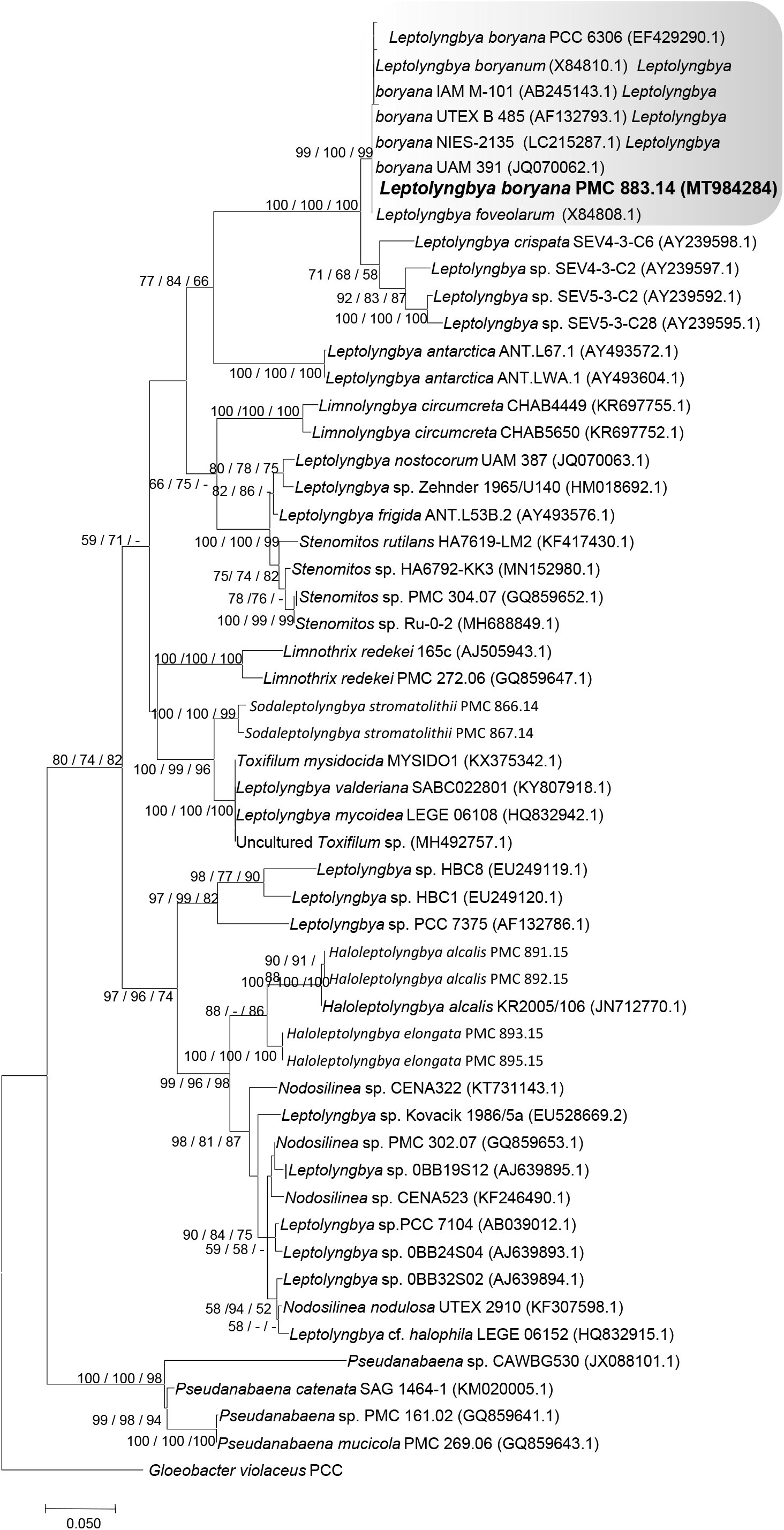
Consensus phylogenetic tree (Maximum likelihood tree presented) based on 16S rRNA gene sequences (54 sequences, 1333 aligned nucleotide position, GTR+G+I model) of representative cyanobacteria strains belonging to the orders Synechococcales, Thermes de Balaruc-Les-Bains’s strain is indicated in bold (with brown color), other species were obtained from Genbank. *Gloeobacter violaceus* PCC 7421 was used as an outgroup. Numbers above branches indicate bootstrap support (>50%) from 1000 replicates. Bootstrap values are given in the following order: maximum likelihood / neighbor joining / maximum parsimony.

The morphological features of the *Leptolyngbya boryana* PMC 883.14 strain were very similar to other *Leptolyngbya* species belonging to the same phylogenetic cluster (Table S3) which made it difficult to distinguish. In Komárek and Anagnostidis (2005), *Leptolyngbya foveolarum* was described with the following characteristics: variously curved filaments, sometimes straight and parallel or tangled, rarely pseudo-branched; trichomes pale to bright blue-green1-2 µm wide; isodiametric cells, rarely longer than wide 0.8-2.2 x 2.5 µm. *Leptolyngbya boryana* was described as follows: curved densely tangled filaments, sometimes pseudo-branched, pale blue-green to colorless, 1.3-2 (3) µm wide, cells more or less isodiametric, somewhat shorter or longer than wide in the main filament 0.6-2 (2.5) x 1.3-3 µm. In other phylogenetic studies, several *Leptolyngbya* species such as *L. foveolarum, L. tenerrima*, and *L. angustata* were considered as synonyms of *L. boryana* (Cuzman et al. 2010). As the genus *Leptolyngbya* was defined based on strains with very thin and simple trichomes (0.5-3.5 μm wide), sheaths present or facultative, without gas vesicles and peripherally arranged thylakoids (Komárek and Anagnostidis 2005; Komárek and Johansen 2015), these morphological features were deemed not sufficiently discriminant to ensure a proper identification; therefore, we consider the strain PMC 883.14 as *Leptolyngbya boryana* based on the former description of this species within the *Leptolyngbya* 16S rRNA gene sequences cluster.

### Characterization of strains belonging to the Oscillatoriales

Among the nine studied strains, four strains belonged to the Oscillatoriales: *Planktothricoides raciborskii* (PMC 877.14), *Laspinema sp*. (PMC 878.14), *Microcoleus vaginatus* (PMC 879.14) and *Lyngbya martensiana* (PMC 880.14).

#### *Planktothricoides raciborskii* (Wołosz) (Suda et M. M. Watanabe, 2002)

*Planktothricoides raciborskii* (PMC 877.14) was isolated from mats covering the mud of the basin (Fig. 1, Table 1). Observations of field samples showed pale-green to brown diffluent thalli (Fig. 10). Trichomes were solitary, pale green to brown, isopolar or heteropolar, cylindrical, variously elongated (100-450 µm). Sheaths were seen occasionally, and were very thin and colorless, with limited motility (gliding).

**Figure 9.**
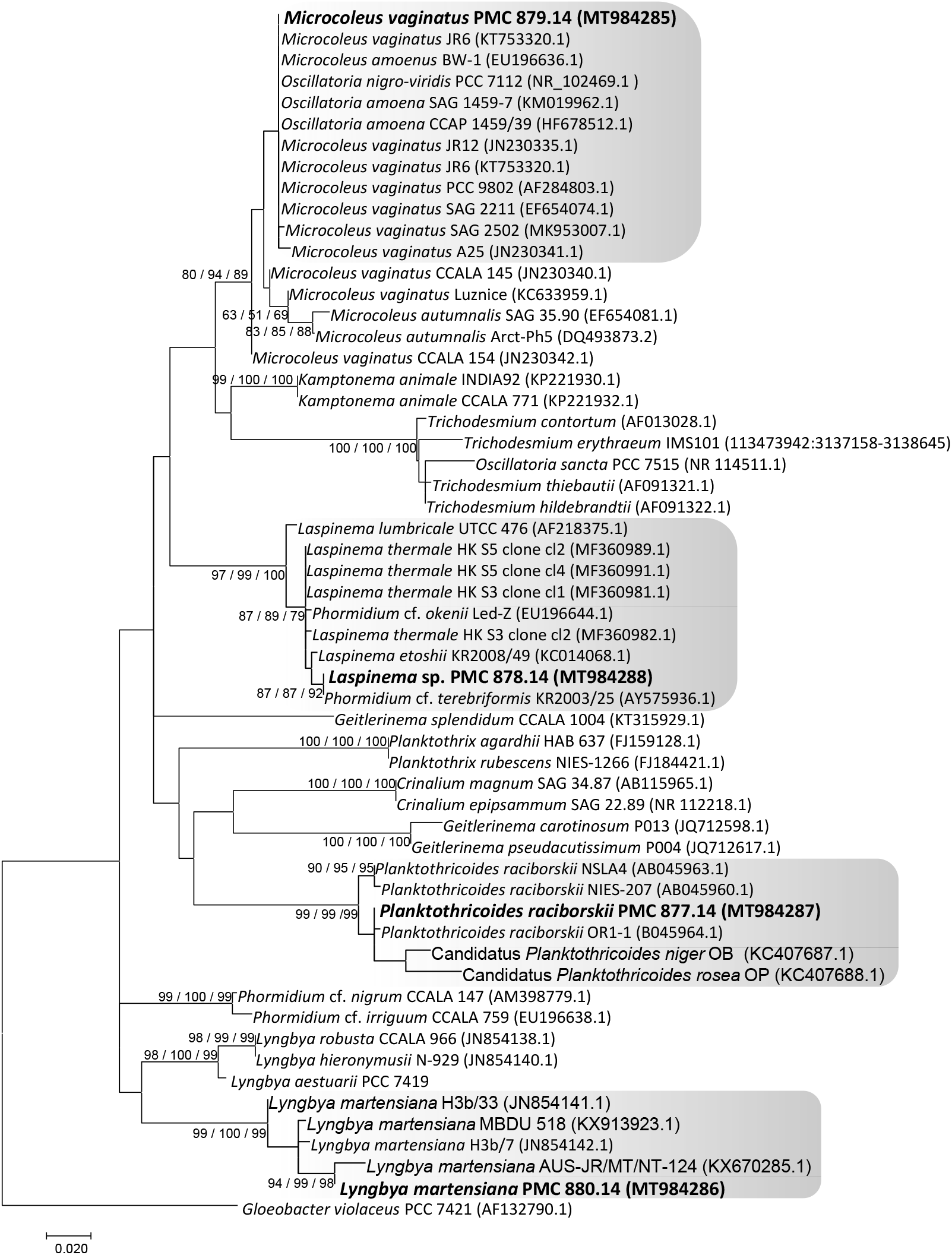
Consensus phylogenetic tree (Maximum likelihood tree presented) based on 16S rRNA gene sequences (57 sequences, 1305 aligned nucleotide positions, GTR+G+I model) of representative cyanobacteria strains belonging to the orders Oscillatoriales, Thermes de Balaruc-Les-Bains’s strains are indicated in bold (colored in blue), other species were obtained from Genbank. *Gloeobacter violaceus* PCC 7421 was used as an out-group. Numbers above branches indicate bootstrap support (>50%) from 1000 replicates. Bootstrap values are given in the following order: maximum likelihood / neighbor joining / maximum parsimony.

**Figure 10.**
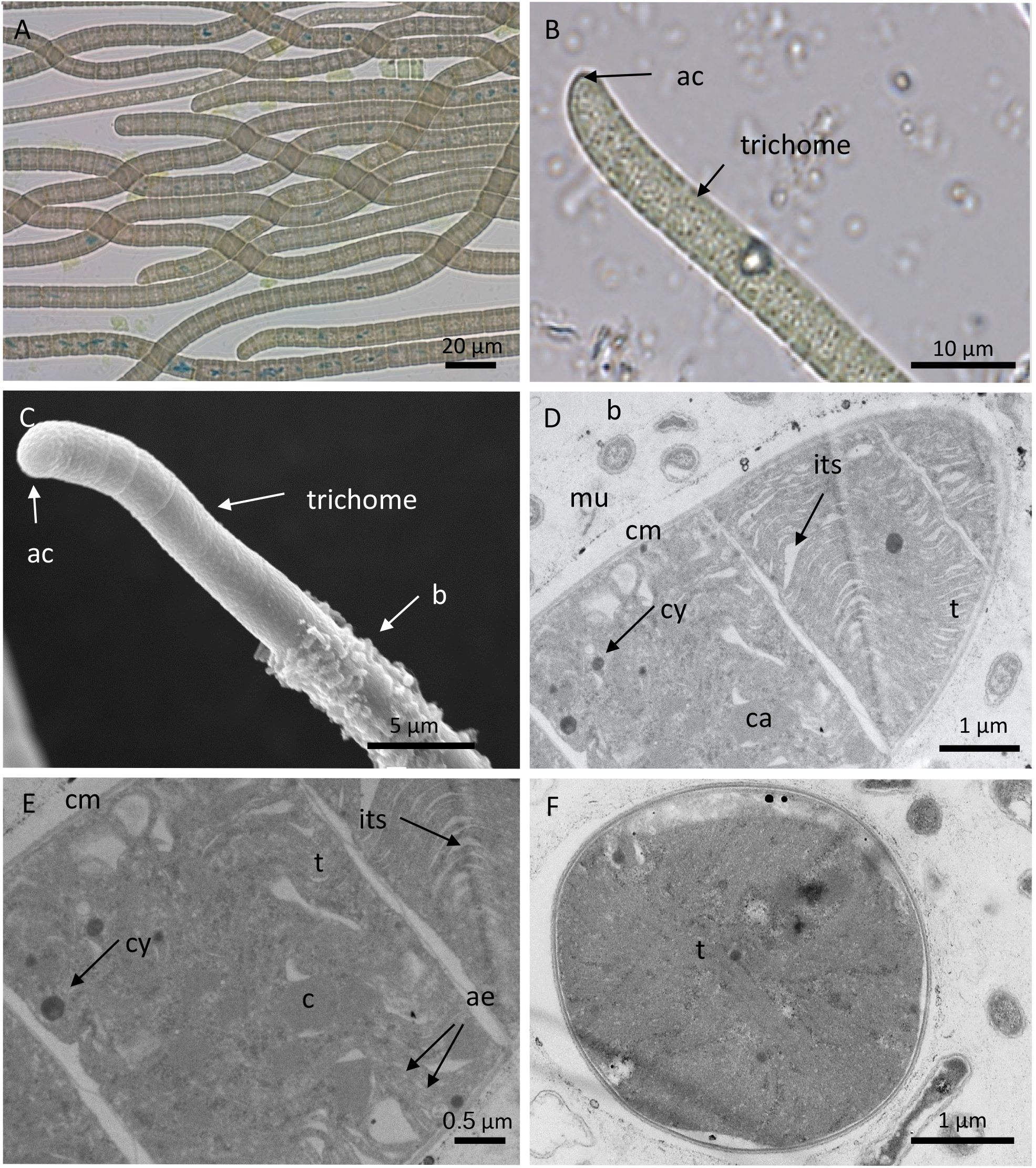
Light microscopy (A, B), SEM (C) and TEM (D-F) micrographs of *Planktothricoides raciborskii* (PMC 877.14) from Thermes de Balaruc-Les-Bains. A: Light microscopy of trichomes from field sample. B: Light microscopy of trichomes in culture. C: trichomes associated or not with bacteria. Longitudinal (C-E) and cross sections (F) of the trichomes showing the ultrastructural details. Abbreviations: ac: apical cell, ae: aerotopes, bact: bacteria, c: carboxysome, cm: cytoplasmic membrane, cy: cyanophycin granules, its: inter thylakoidal space, r: reserves (lipids, polyphosphates, poly-β-hydroxybutyrate), t: thylakoids, vs: clear vesicle.

Trichomes were attenuated towards extremities, constricted or no constricted and sometimes slightly bent near the apex (Fig. 10). Cells were cylindrical, shorter than wide, 5-7.3 µm wide and 3.3-5.4 µm long, with a length:width ratio of 0.7. Apical cells were rounded, conical, more or less tapered, sometimes bent but not sharply pointed, without calyptra (Fig. 10; Table S4). Cell content was granular with cyanophycin granules and carboxysomes (polyhedral or dense bodies; Fig. 10). These proteinaceous micro-compartments play a major role in the carbon-fixation process by increasing the CO_2_ concentration around RuBisCO, thus promoting carboxylation at the expense of oxygenation. Sheaths were generally thin and some trichomes were associated with bacteria under our laboratory culture conditions. Cross-walls were clearly visible with TEM (Fig. 10) and the thylakoids were arranged radially in cross sections of the trichomes.

The 16S rRNA gene sequence comparisons (BLAST) showed that the PMC 877.14 strain shared ≥ 99% sequence identity with other *Planktothricoides* strains (data not shown). The phylogenetic tree of the sequences of *Planktothricoides* produced two main clusters (Fig. 9). The first cluster was primarily composed of strains isolated from freshwater habitats, the one from Thermes de Balaruc-Les-Bains, one strain from Thailand, one from Australia, one from Japan and one from Singapore. The second cluster grouped the strains *Candidatus ‘*Planktothricoides niger*’* and *Candidatus ‘*Planktothricoides rosea’ from periphyton mats of tropical marine mangroves in Guadeloupe (Guidi-Rontani et al. 2014). The two species, *Candidatus* ‘Planktothricoides niger*’* and *Candidatus* ‘Planktothricoides rosea’, have been distinguished by their phycoerythrin pigment content. The *Planktothricoides* strain from Thermes de Balaruc-Les-Bains (PMC 877.14) contained two dominant phycobilin pigments, phycoerythrin (PE) and phycocyanin (PC), with a PC:PE ratio = 1 (Fig. S1, Table S4). As the color of trichomes differs between field samples and laboratory cultures, we suggest that this strain can either undergo complementary chromatic adaptation (Grossman, 2003) or modify its PC:PE ratio depending on light wavelength or penetration through the biomass. This characteristic is clearly different from the type description of *Planktothricoides raciborskii* (Wołosz.) Suda and M. M. Watanabe (Suda et al., 2002; Komárek et al. 2004), where phycoerythrin was described as absent and complementary chromatic adaptation was not observed. The presence of PE should not be considered as sufficient to assign the strain PMC 877.14 to either *Planktothricoides niger* or *Planktothricoides rosea*, and we considered the phylogenetic data strong enough to label the PMC 877.14 strain as *Planktothricoides raciborskii*.

#### *Laspinema* sp (F. Heidari & T. Hauer, 2018)

The *Laspinema* strain (PMC 878.14) was isolated from the mud covering the maturation basin of Thermes de Balaruc-Les-Bains. Examination of field samples showed a pale green to brown diffluent thallus (Fig. 11). In culture, the *s*train showed green or blackish green thallus formed mats and its trichomes were bright blue-green often gradually attenuated at the ends, long, straight, not constricted at the finely granulated cross-walls, bent, hooked and intensely motile (Fig. 11). Cells were always shorter than wide, 3.8-4.8 µm wide, 2.3-3.3 µm long with a length:width ratio of 0.7. Sheaths were generally absent (Fig. 11). The apical cells were obtusely-rounded to rounded-conical without calyptra, and often with thickened outer cell walls (Fig. 11).

**Figure 11.**
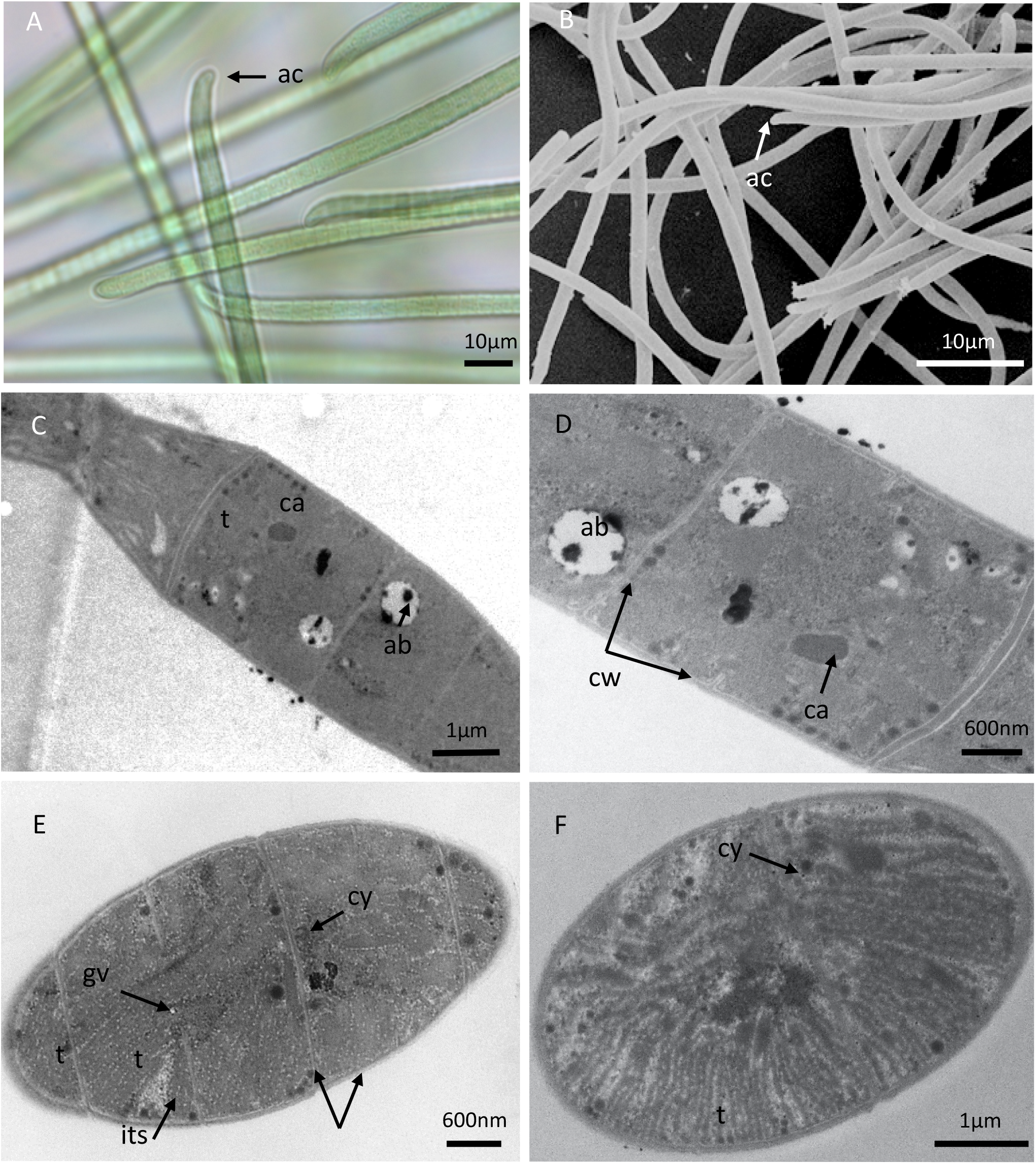
Light microscopy (A), SEM (B) and TEM (C-D-E-F) micrographs of *Laspinema sp*. (PMC 878.14) from Thermes de Balaruc-Les-Bains. A: filaments with typical shape of apical cell. B: observations of filaments. Longitudinal (C-E) and cross sections (F) of trichomes showing the ultrastructural details. Abbreviations: ab: beta-hydroxybutyric acid, ac: apical cell, ca: carboxysome, cy: cyanophycin granule, gv: gas vesicles, its: inter thylakoidal space, t: thylakoids.

The thylakoids visible in cross sections of the trichomes were radially arranged with several interthylakoidal spaces (Fig. 11). Other components such as numerous small gas vesicles or aerotopes, which promote flotation of the filaments, carboxysomes, cyanophycin granules (nitrogen reserves), glycogen, lipids, or poly-β-hydroxybutyrate acting as carbon and energy sources, and polyphosphate granules (phosphorus reserves) were observed within the cells

The 16S rRNA gene sequence analysis (BLAST) showed that the PMC 878.14 strain shared ≥99% sequence identity with other *Laspinema* sp. strains (data not shown). The phylogenetic tree of *Laspinema* strains had one robust cluster (Fig. 9) composed of several species including *L. lumbricale, L. thermale* and *L. etoshii*. Comparison of the morphological characteristics of the different species (Table S5) did not allow us to assign a species name to the studied strain. Thus, strain PMC 878.14 was named *Laspinema* sp. based on morphology (Table S5) and phylogenetics (Fig. 9).

#### *Microcoleus vaginatus* (Gomont ex Gomont 1892)

*Microcoleus vaginatus* (PMC 879.14) was isolated from the wall of the retention basin. The field samples showed a dense green or blackish green thallus (Fig. 12). In culture, the thallus was green or blackish green, forming mats and showing motility (gliding). Trichomes were often gradually attenuated at the ends, with no constrictions at the cross-walls (Fig. 12). The cell wall envelope consisted of a cytoplasmic membrane bounded by relatively thin wall layers. The apical cells were rounded with rounded or truncated calyptra, often with thickened outer cell walls (Fig. 12). Cells were always shorter than wide, 4.7-6.2 µm wide by 2.4-4.8 µm long, with a length:width ratio of 0.7 (Table S6).The thylakoids formed fascicle-like aggregations arranged irregularly in the cells. The cells contained several large cyanophycin granules, some carboxysomes and numerous small gas vesicles or aerotopes (Fig. 12).

**Figure 12.**
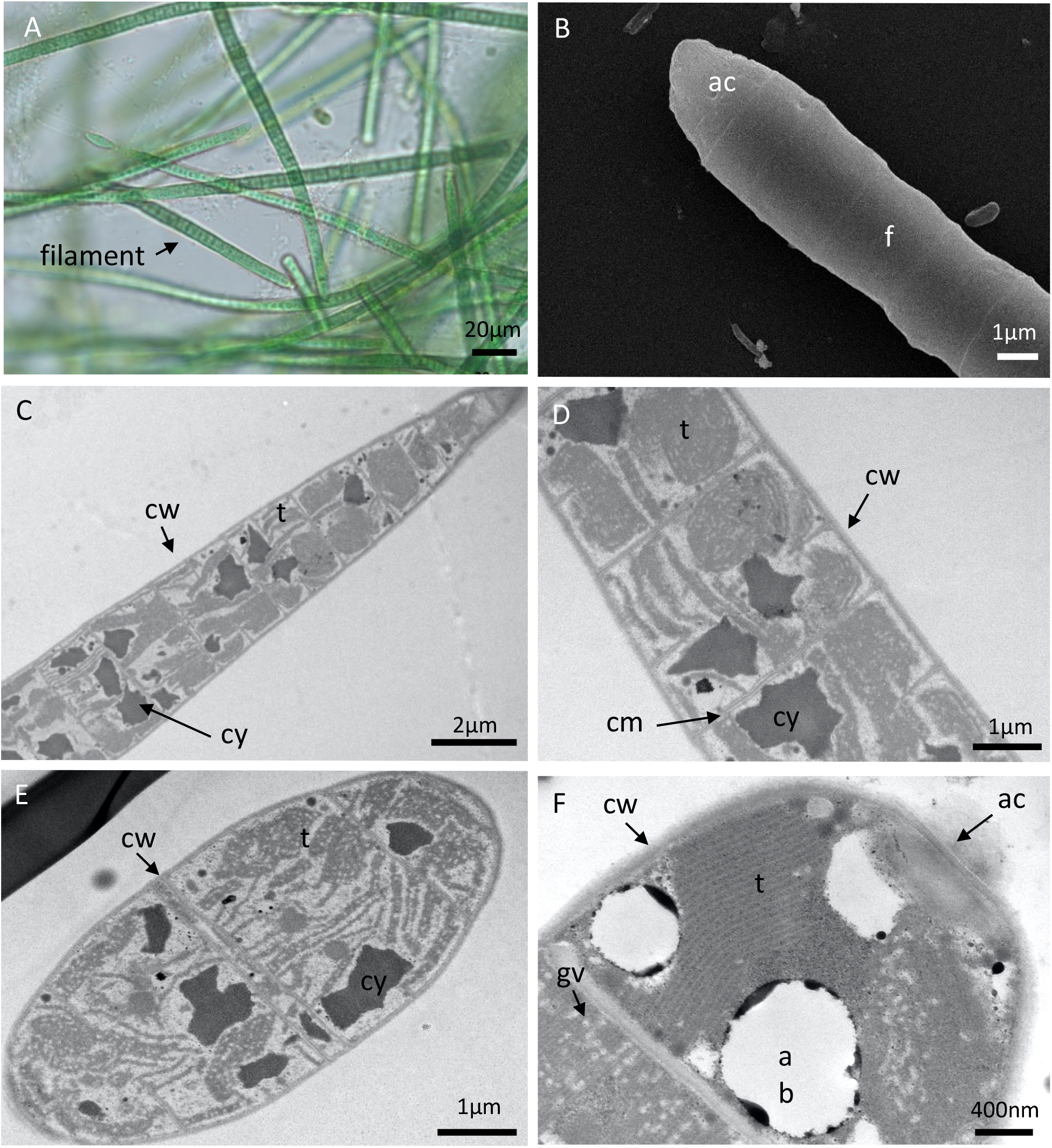
Light microscopy (A), SEM (B) and TEM (C-D-E-F) micrographs of *Microcoleus vaginatus* (PMC 879.14) from Thermes de Balaruc-Les-Bains. A: entangled filaments with sheath. B: filaments with apical cell. Longitudinal (C-D-E-F) of trichomes with sheaths showing the ultrastructural details. Abbreviations: ab: acid beta-hydroxybutyric, ac: apical cell, cy: cyanophycin granule, f: filament, t: thylakoids.

The 16S rRNA gene sequence analysis (BLAST) showed that PMC 879.14 shared ≥98% sequence identity with other *Microcoleus vaginatus* strains (data not shown). The phylogenetic tree placed the sequence in a well-grouped cluster containing numerous *M. vaginatus* sequences (Fig. 9). Some sequences belonging to *Oscillatoria amoena* and *O. nigro-viridis* were also grouped in this cluster, probably because they were assigned before the taxonomic revision of this group (Strunecky et al. 2013). According to phylogeny and following the recent taxonomic revision, the strain PMC 879.14 was assigned to *Microcoleus vaginatus*.

#### *Lyngbya martensiana* (Meneghini ex Gomont 1892)

*Lyngbya martensiana* (PMC 880.14) was isolated from the epilithic biofilm of the retention basin of Thermes de Balaruc-Les-Bains. Field examinations and laboratory cultures showed bright-green thalli composed of tangled filaments, usually arranged in parallel (Fig. 13). Filaments were long, straight or variously flexuous with thick colorless sheaths. Trichomes were cylindrical and not constricted at the cross-walls. The granular cell contents were pale blue-green. Cells were discoid, shorter than wide, 4.8-6.6 (5.3±0.47) µm wide, 1.2-2.1 (1.6±0.2) µm long with a length:width ratio of 0.3. Apical cells were widely rounded, hemispherical or depressed-hemispherical without calyptra (Fig. 13). The thylakoids formed fascicle-like aggregations arranged irregularly in the cells and situated near walls and cross-walls.

**Figure 13.**
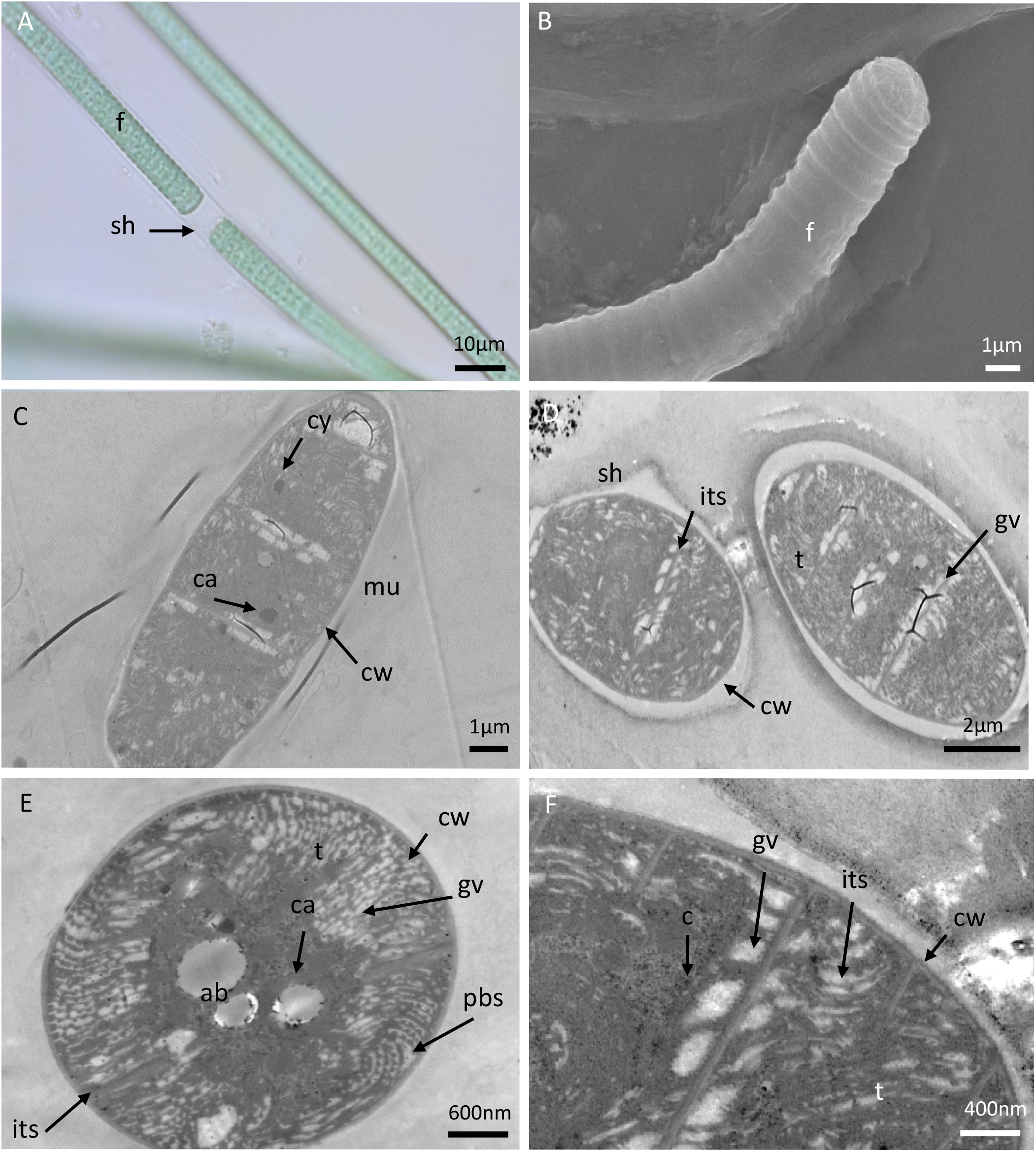
Light microscopy (A), SEM (B) and TEM (C-D-E-F) micrographs of *Lyngbya martensiana* (PMC 880.14) from Thermes de Balaruc-Les-Bains. A: filaments with sheath. B: observation of a filament. Longitudinal section (C-D-F) and cross section showing the ultrastructural details. Abbreviations: ab: beta-hydroxybutyric acid, ca: carboxysome, cy: cyanophycin, cw: cell wall, its: inter thylakoidal space, f: filament, pbs: phycobilisomes (photosynthetic antenna), s: sheath, t: thylakoids.

Numerous pores penetrating the cross-walls between the cells were clearly visible (Fig. 13). Gas vesicles occurred throughout the cells, which also contained reserve granules and some carboxysomes (Fig. 13). The 16S rRNA gene sequence analysis showed that PMC 880.14 shared ≥96% sequence identity with other *Lyngbya martensiana* strains (data not shown). The phylogenetic tree of the sequences of *Lyngbya martensiana* strains produced one well-grouped cluster (Fig. 9). The morphological characteristics (Table S7) and the well-grouped *Lyngbya martensiana* cluster allowed us to name the PMC 880.14 strain as *Lyngbya martensiana* (Gomont M. 1892).

### Characterization of strains belonging to the Order Nostocales

Among the nine studied strains, three belonged to the Nostocales: *Nosto*c sp. (PMC 881.14), *Aliinostoc* sp. (PMC 882.14) and *Calothrix* sp (PMC 884.14).

#### *Nostoc* sp (Vaucher ex Bornet & Flahault, 1886)

*Nostoc* sp. (PMC 881.14) was isolated from the epilithic biofilm on the wall of the retention basin. Field samples had a dense thallus with blackish green or brownish macroscopic colonies (Fig. 15). These macroscopic colonies resulted from several small, amorphous, fine, thin and mucilaginous colonies. In culture, old colonies (up to 130 µm in diameter) were enveloped by a firm periderm that could break and release young colonies. Within the colonies, the filaments were very densely tangled, flexuous and/or coiled. They were blackish green or brownish, ∼5 µm wide, highly constricted at cross-walls, uniseriate, unbranched, isopolar, moniliform and/or curled (Fig. 15). Cells were isodiametric, well separated from one another by cross-walls, wider than long, 2.8-5.5 (4.5±0.9) μm wide, 2.6-4.6 (3.4±0.5) µm length with a length:width ratio of 0.7. The apical cells were spherical to conical (data not shown). The heterocytes were 2.6-5.7 (3.4±0.6) µm wide and 2.3-5.1 (3.8±0.7) µm long. Akinetes were never observed during the three years of culture, even when the strain was cultivated in Z8X medium without a nitrogen source to induce stress conditions. An extensive gelatinous matrix was observed resulting in a long internal diffusion path from the surface to the trichomes. A sheath with a periplasmic membrane was clearly visible by TEM (Fig. 15). The thylakoids were fascicular with irregular spherical formations (Fig. 15).

**Figure 14.**
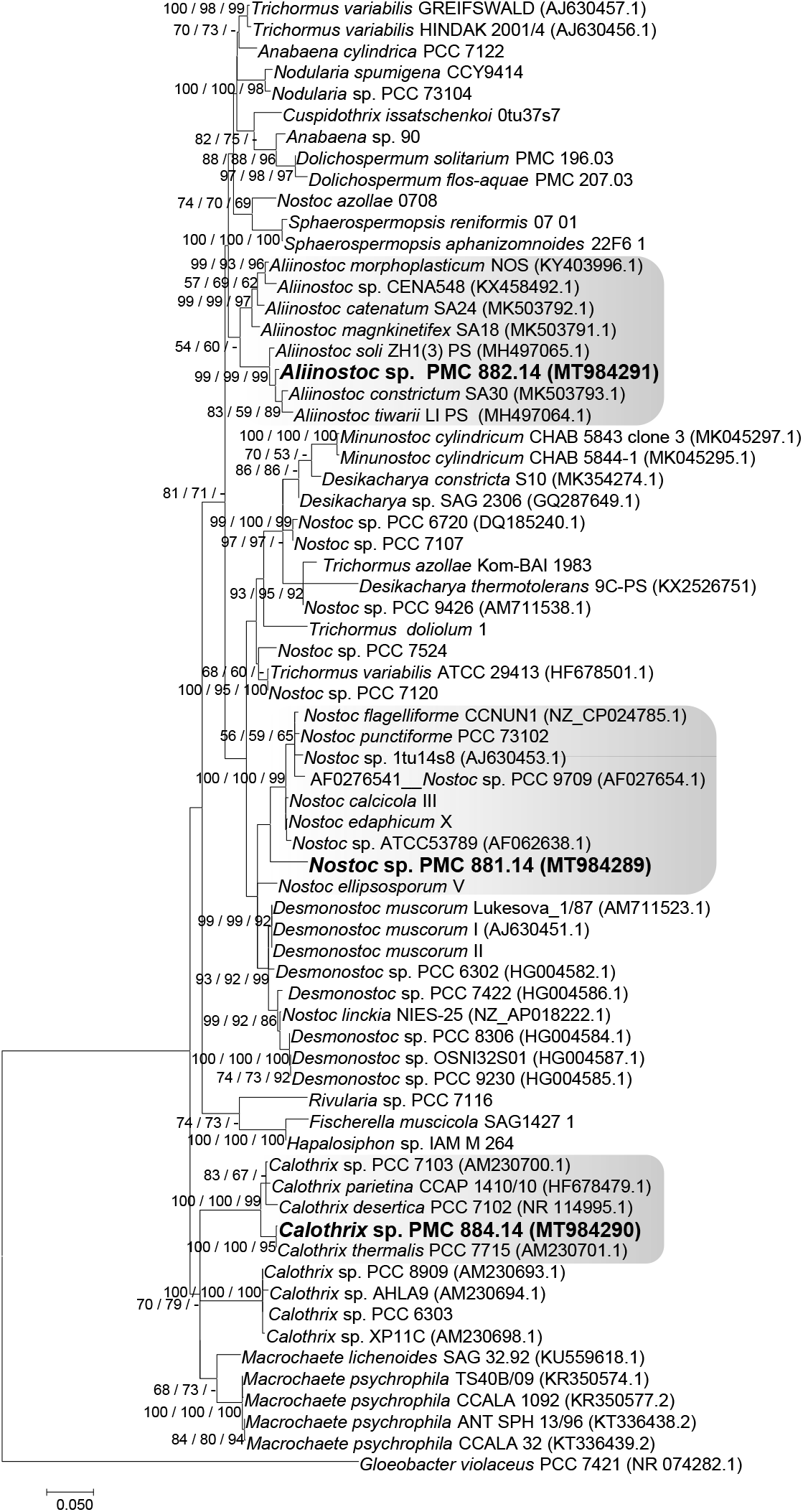
Consensus phylogenetic tree (Maximum likelihood tree presented) based on 16S rRNA gene sequences (69 sequences, 1310 aligned nucleotide position, GTR+G+I model) of representative cyanobacteria sequences (Genbank) belonging to the Nostocales and the studied strains from Thermes de Balaruc-Les-Bains are indicated in bold (with purple color). *Gloeobacter violaceus* PCC 7421 was used as an out-group. Numbers above branches indicate bootstrap support (>50%) from 1000 replicates. Bootstrap values are given in the following order: maximum likelihood / neighbor joining / maximum parsimony.

**Figure 15.**
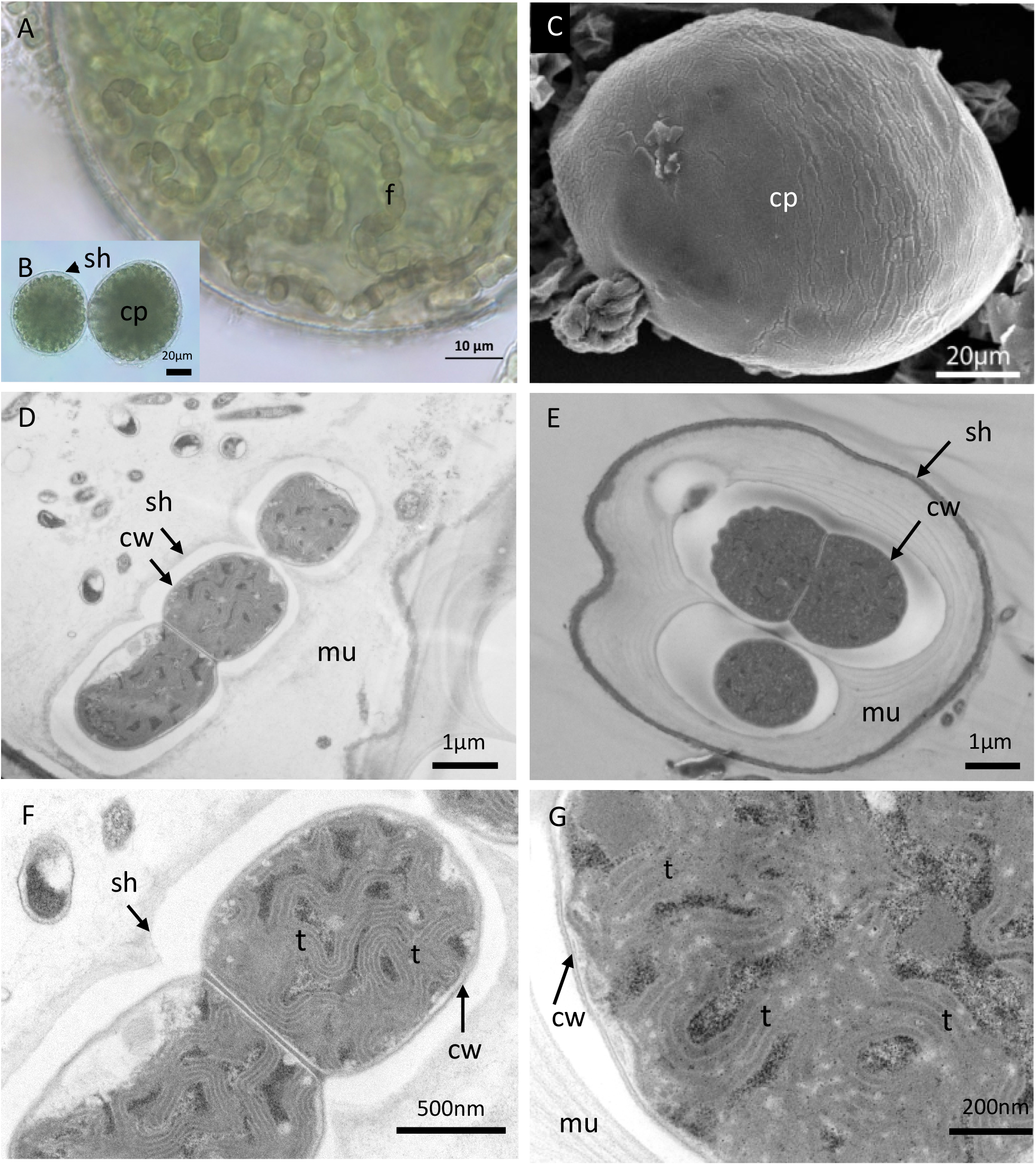
Light microscopy (A-B), SEM (C) and TEM (D-E-F-G) micrographs of *Nostoc* sp. (PMC 881.14) from Thermes de Balaruc-Les-Bains. A: Observation of the filaments located inside the capsule, B: Observation of two colonies, spherical, irregular in their shape, dull olive to brownish-colored and gelatinous. Filaments very densely entangled, flexuous, coiled inside the colonies. C: external observation of a capsule. Longitudinal (D-F-G) and cross sections (E) showing the ultrastructural details. Abbreviations: cp: capsule, cw: cell wall, f: filament, mu: mucilage, sh: sheath, t: thylakoids.

The 16S rRNA gene sequence analysis showed that PMC 881.14 strain shared ≥ 96% sequence identity with other *Nostoc* sp. strains (data not shown). The phylogenetic tree based on 16S rRNA gene sequences showed that the PMC 881.14 strain was grouped into a unique *Nostoc* cluster (Fig. 14) with a high bootstrap value. Within this cluster three sequences were not assigned at the species level, and the five others were assigned to different species: *N. calcicola, N. punctiforme, N. flagelliforme, N. edaphicum, N. ellisporum*.

It is now well known that *Nostoc* is a complex genus. The species belonging to *Nostoc* were generally described as microscopic, spherical, oval, ovoid, irregular in their colony form, gelatinous, irregularly clustered, amorphous, dull olive to brownish gelatinous clusters free-living on mud (Table S8). Because of this heterogeneity, it is difficult to clearly differentiate related taxa within the genus based solely on morphology (Singh et al. 2016 and 2020). Previous studies using a polyphasic approach led to the splitting of *Nostoc* into several genera, including *Mojavia, Desmonostoc* and *Compactonostoc* (Hrouzek et al. 2003; Řeháková et al. 2007, Cai et al, 2019). It is difficult to separate these groups within *Nostoc sensu lato* based only on morphological characteristics (Table S8). Following the concept that a taxonomic unit (family, genus, species) must be included in only one phylogenetic lineage (Komárek, 2018), the strain PMC 881.14 was considered as a *Nostoc* sp. because our results did not allow us to give any further species assignment to this strain.

#### *Aliinostoc* sp (S.N. Bagchi, N. Dubey & P. Singh, 2017)

*Aliinostoc* sp. (PMC 882.14) was isolated from the epilithic biofilm on the wall of the retention basin. Field samples showed free blue-green filaments (Fig. 16), while in culture the filaments were flexuous and entangled. The trichomes were pale blue-green, cylindrical, thin (< 6 µm wide), and highly constricted at cross-walls (Fig. 16). Cells were well separated from each other by cross-walls, isodiametric, and longer than wide, 3.4-5.6 (4.5±0.6) μm wide by 2.6-6.4 (4.6±1.0) µm long with a length:width ratio of 1.0 (Table S9). SEM showed that apical cells were conical (Fig. 16). The heterocytes were 4.1-6.5 (5.4±0.5) µm wide by 4.1-6.8 (5.4±0.6) µm length and akinetes were 4.5-7.4 (5.9±0.8) µm wide by 4.5-9.0 (7.2±1.0) µm long (Table S9). No sheath was visible under TEM (Fig. 16). The thylakoids were fascicular with irregular spherical formations (Fig. 16). The cells contained several reserve granules and numerous carboxysomes (Fig. 16). The heterocyst TEM observations showed a few thylakoids and large cyanophycin granules.

**Figure 16.**
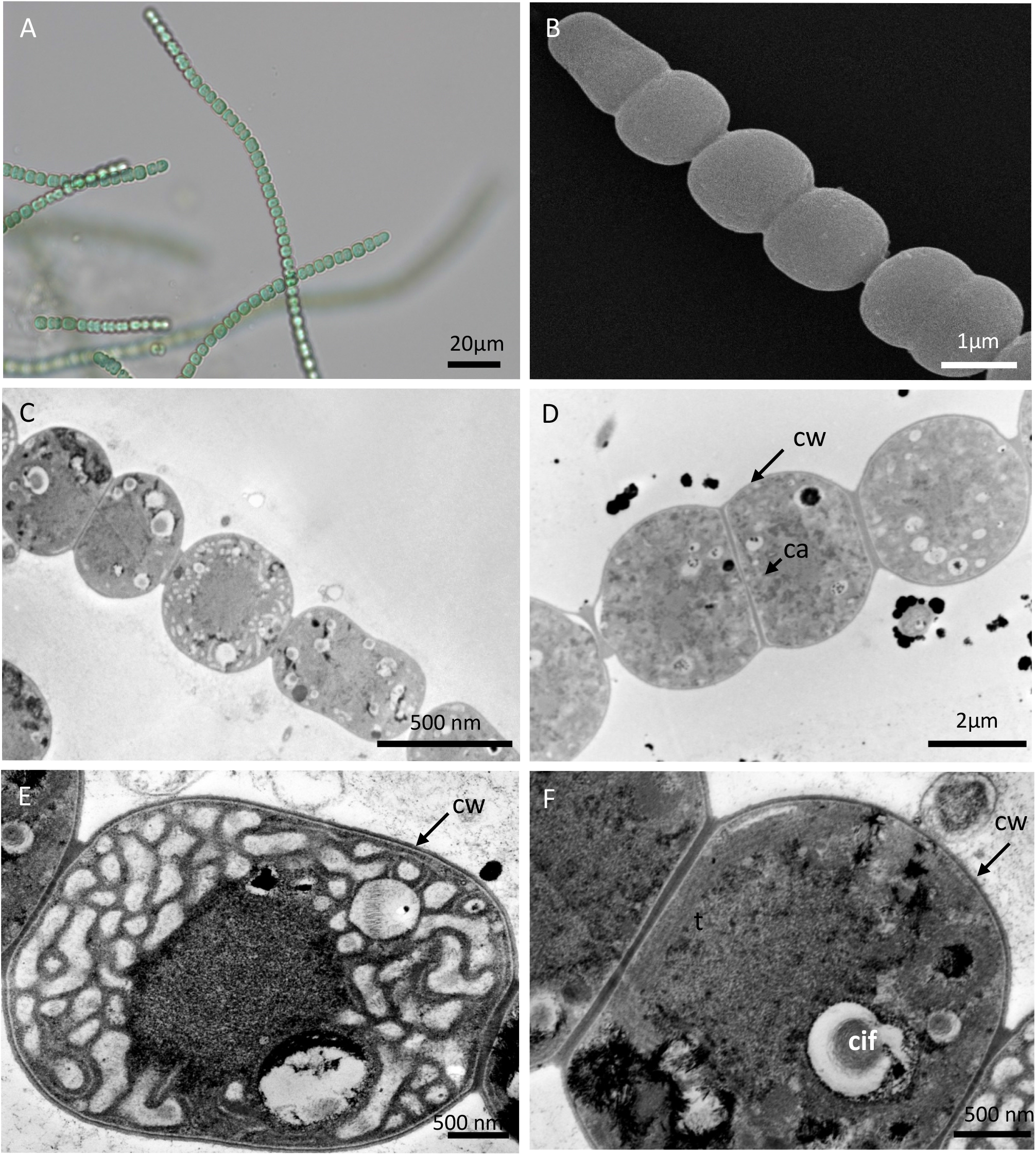
Light microscopy (A), SEM (B) and TEM (C-F) micrographs of *Aliinostoc* sp. (PMC 882.14) from Thermes de Balaruc-Les-Bains. A: Light microscopy of filament from field sample. B: SEM observations of trichome. Longitudinal sections(C-F) of an heterocyte (E) and vegetative cells (C, D, F) showing the ultrastructural details. Abbreviations: ca: carboxysome, cif: carboxysome in formation, cw: cell wall, its: inter thylakoidal space, t: thylakoids.

The 16S rRNA gene sequence analysis showed that PMC 882.14 shared ≥ 98% sequence identity with other *Aliinostoc* sp. Strains (data not shown). The trees based on 16S rRNA gene sequences placed PMC 882.14 within a well-grouped cluster of *Aliinostoc* sequences (Fig. 14). Within this cluster, several sequences were identified as *Aliinostoc tiwarii, A. constricutum, A. soli, A. magnkinetifex, A. catenatum* and *A. morphoplasticum*, mainly based on morphological characteristics (Table S9). *Aliinostoc* sp. Showed close morphological resemblance to the genera *Nostoc* and *Trichormus*. A diacritical feature for *Aliinostoc* is the presence of motile hormogonia with gas vesicles (Bagchi et al. 2017). The authors also stated that filaments must be loosely arranged with variable tendencies for coiling. These organisms were generally isolated from habitats rich in dissolved ions and salts, with high water conductivity. As for the *Nostoc* sp. PMC 881.14, description and morphology alone were of little help in assigning the strain to *Aliinostoc sensu lato* (Bagchi et al. 2017) (Table S9). Based on the rule that a taxonomic unit (i.e. genus, species) must be included in only one phylogenetic lineage (Komárek, 2018), the strain PMC 882.14 was named *Aliinostoc* sp. because our data were not sufficient to give a species assignment.

#### *Calothrix* sp (C. Agardh ex Bornet & Flahaut, 1886)

*Calothrix* sp. (PMC 884.14) was isolated from the biofilm at the surface of the mud of the retention basin. Observations of field samples showed a brown filamentous thallus basally attached to the substratum and forming thin mats. In culture, the filamentous thallus formed small brownish-green tufts (Fig. 17). Filaments were aggregated, flexuous, heteropolar, widened near the base and narrowed towards the ends, without false branching. The terminal cells were of variable length, conical, bluntly pointed and tapered. The sheath was generally firm and colorless. The trichomes were of different lengths (> 800 µm long), pale blue-green, olive-green to greyish-green, more or less constricted at the cross-walls, and narrowed at the apex: 6.7-10.4 (8.1±0.7) µm wide at the base, 4.7-7.7 (5.8±0.6) µm in the middle and 3.5-5 (4.4±0.4) µm near the ends (Fig. 17, Table S10). Vegetative cells were more or less quadratic, generally shorter than wide in the upper part of the trichome and/or longer than wide in the lower part of the trichome, 5.2-7.9 (6.6±0.7) μm wide by 4.8-6.2 (5.5±0.4) µm long. Heterocytes were basal, single, hemispherical, 4.7-8.2 (6.2±0.9) μm wide by 3.5-7.3 (4.9±0.9) µm long. Akinetes were not observed during the three years the cultures were maintained, even when strain PMC 884.14 was cultivated under stress conditions.

**Figure 17.**
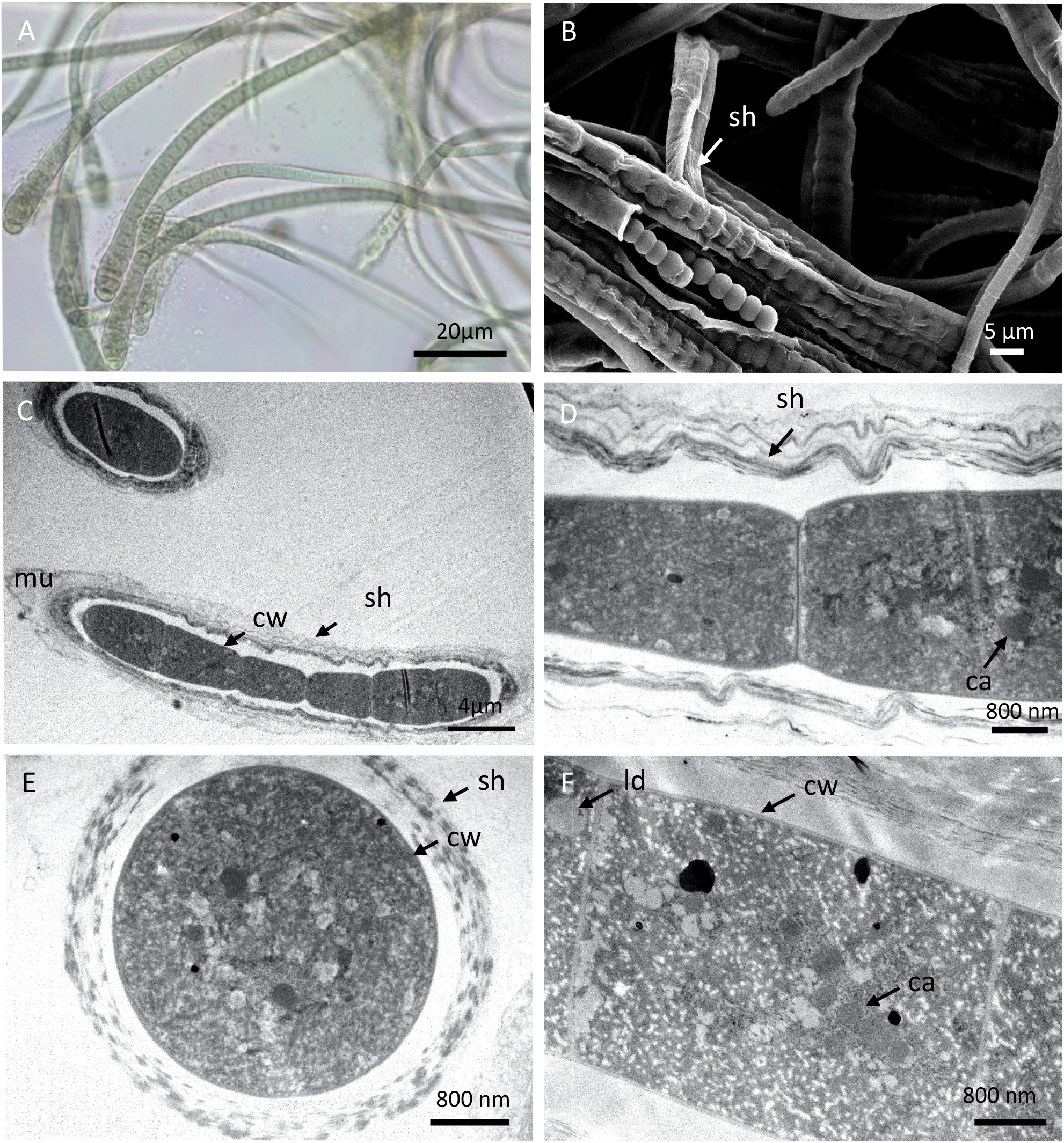
Light microscopy (A), SEM (B) and TEM (C-D-E-F) micrographs of *Calothrix sp*. (PMC 884.14) from Thermes de Balaruc-Les-Bains. A: entangled filaments with sheath. B: observations of filament. Longitudinal (C-D-F) and cross sections (E) of trichomes with sheaths showing the ultrastructural details. Abbreviations: ca: carboxysome, ld: lipid droplet, cw: cell wall, sh: sheath, t: thylakoids, mu: mucilage.

The 16S rRNA gene sequence analysis showed that PMC 884.14 shared ≥ 99% sequence identity with other *Calothrix* sp. strains (data not shown). Phylogenetic trees grouped PMC 884.14 into a coherent *Calothrix* clade (Fig. 14) in which the sequences were mainly assigned to *Calothrix* sp. Recent revision suggested that the genus *Calothrix* should be split into several genera including *Tolypothrix* and *Rivularia* (Berrendero et al. 2016). Based on a polyphasic study of 135 heterocytous tapering cyanobacteria, of which 99 were identified as *Calothrix*, these authors showed that the majority of *Calothrix* sequences were spread among four lineages, two of which were marine, while the other two were freshwater and soil strains (Berrendero et al. 2011, 2016). As the sequences for the *Calothrix* type species (*C. confervicola*) were lacking, and no culture was available in any collection, we could not say which *Calothrix* clade represents the true *Calothrix*. It would be helpful to conduct more phylogenetic analyses to elucidate the phylogeny of the heteropolar, tapering taxa within Nostocales. Based on the rule that a taxonomic unit must be included in only one phylogenetic lineage (Komárek, 2018), PMC 884.14 was assigned to *Calothrix* sp. as this study did not allow us to give a species name to this strain.

### Potential therapeutic properties of cyanobacterial strains from the muds of the thermal springs of Balaruc-les-Bains

A polyphasic approach is still the best way to correctly determine the taxonomy of a new isolate (Komárek 2018, 2020). Using morphological, ultrastructural and molecular methods, the nine cyanobacterial isolates from the muds of Thermes de Balaruc-Les-Bains, were clearly identified as belonging to the orders Chroococcales: *Pseudo-chroococcus couteii*; Synechococcales: *Leptolyngbya boryana*; Oscillatoriales: *Planktothricoides raciborskii, Laspinema* sp., *Microcoleus vaginatus, Lyngbya martensiana* and Nostocales: *Nostoc* sp., *Aliinostoc* sp., *Calothrix* sp. This taxonomic diversity along with literature reports of the bioactive metabolite synthesis potential of these taxa allowed us to hypothesize that some of the metabolites produced by these strains may be active in the mud therapy at Thermes de Balaruc-Les-Bains. A recent literature review (Demay et al. 2019) emphasized this great potential (Table S11). All cyanobacteria are known to synthesize pigments such as chlorophylls, phycobiliproteins and carotenoids (Fig. S1), which can have antioxidant or anti-inflammatory properties (Table S11). These pigments were synthesized by the studied isolates in different percentages, depending on the strain and the physiology (Table S11). In addition to pigment production, *Lyngbya, Calothrix* and *Chroococcus* strains have attracted further interest because they produce bioactive molecules such as mycosporine-like amino acids (MAAs) and the yellow-brown scytonemin pigments with reported antioxidant, anti-inflammatory and UV-protecting activities. Numerous bioactive compounds from strains belonging to the *Nostoc* genus have also been described, including the anti-inflammatory metabolites aeruginosins and scytonemins. *Planktothricoides, Aliinostoc, Laspinema* sp., *Microcoleus* and the new genus *Pseudo-chroococcus* were not reported in Demay et al. (2019) to have antioxidant and/or anti-inflammatory activities.

Misidentification of Oscillatoriales with their very simple filamentous organization and no diacritical characteristics, occurs frequently and some isolates could have been described as *Phormidium, Lyngbya* or other simple organized Oscillatoriales.

## CONCLUSIONS

Cyanobacterial taxonomy has changed substantially over the last few years thanks to revisions of the morphological criteria and the use of gene sequencing. To resolve the taxonomic discrepancies between morphological and 16S rRNA data and better assign new isolates of cyanobacteria, the use of whole genome sequencing is warranted. Approximately 1440 cyanobacterial genomes (complete or in progress) are currently available (S. Halary, personal communication) in GenBank (NCBI Datasets “cyanobacteria”, accessed on 9 Sept. 2020). However, this growing number of available genomes may hide the low representativeness of the diversity of cyanobacteria since most sequenced strains belong to the orders Nostocales and Synechococcales with 457 and 600 available assemblies, respectively. For example, very few genomes were available for the genera assigned to Balaruc strains with 2-8 genomes available for *Planktothricoides, Lyngbya* and *Microcoleus* and none for *Aliinostoc* and *Pseudo-chroococcus*. Correct assignment of new isolates is essential for understanding the relationships among strains and identifying beneficial therapeutic properties such as those possessed by the mud therapy strains from Thermes de Balaruc-Les-Bains. The isolates maintained in collections will allow in-depth investigation of their potential for production of bioactive metabolites using genomic and biochemical methods.

## AUTHOR CONTRIBUTIONS

SH, CY, CD and CB have worked on the design of the experiments. CD, SH, BP, GT and JD have performed the experiments. CD, SH, BP and CB have performed the data analysis. CD, SH, JD, SD, BM, CY and CB have written and edited the paper.

## ACKNOLEDMENTS

This work was supported by the ANRT through a PhD grant awarded to J. Demay. We would like to thank the UMR 7245 MCAM, Muséum National d’Histoire Naturelle, Paris, France for laboratories facilities and the Thermes de Balaruc-Les-Bains for complimentary founds. The authors thanks Dr P. Méric (Thermes of Balaruc-Les-Bains) for his support during prospecting campaigns on and C. Djediat, A. Marme and S. Jaber for preliminary TEM studies. This work was supported by the “Société Publique Locale d’Exploitation des Thermes de Balaruc-Les-Bains” (SPLETH), the town of Balaruc-Les-Bains and the National Muséum of Natural History. The authors also would like to thank Peerwith (https://www.peerwith.com) for English language editing.

## DEDICATION

This manuscript is dedicated to the memory of late Professor Alain Couté, who made an outstanding contribution to the taxonomy of cyanobacteria and microalgae in the last decades. His work covers a wide range of studies dealing with photosynthetic microorganisms from marine and freshwater habitats, and also from terrestrial and extreme environments. Alain Couté’s contribution will be always relevant and was a driver for shaping this manuscript.

## Supplementary materials

**Figure S1.**
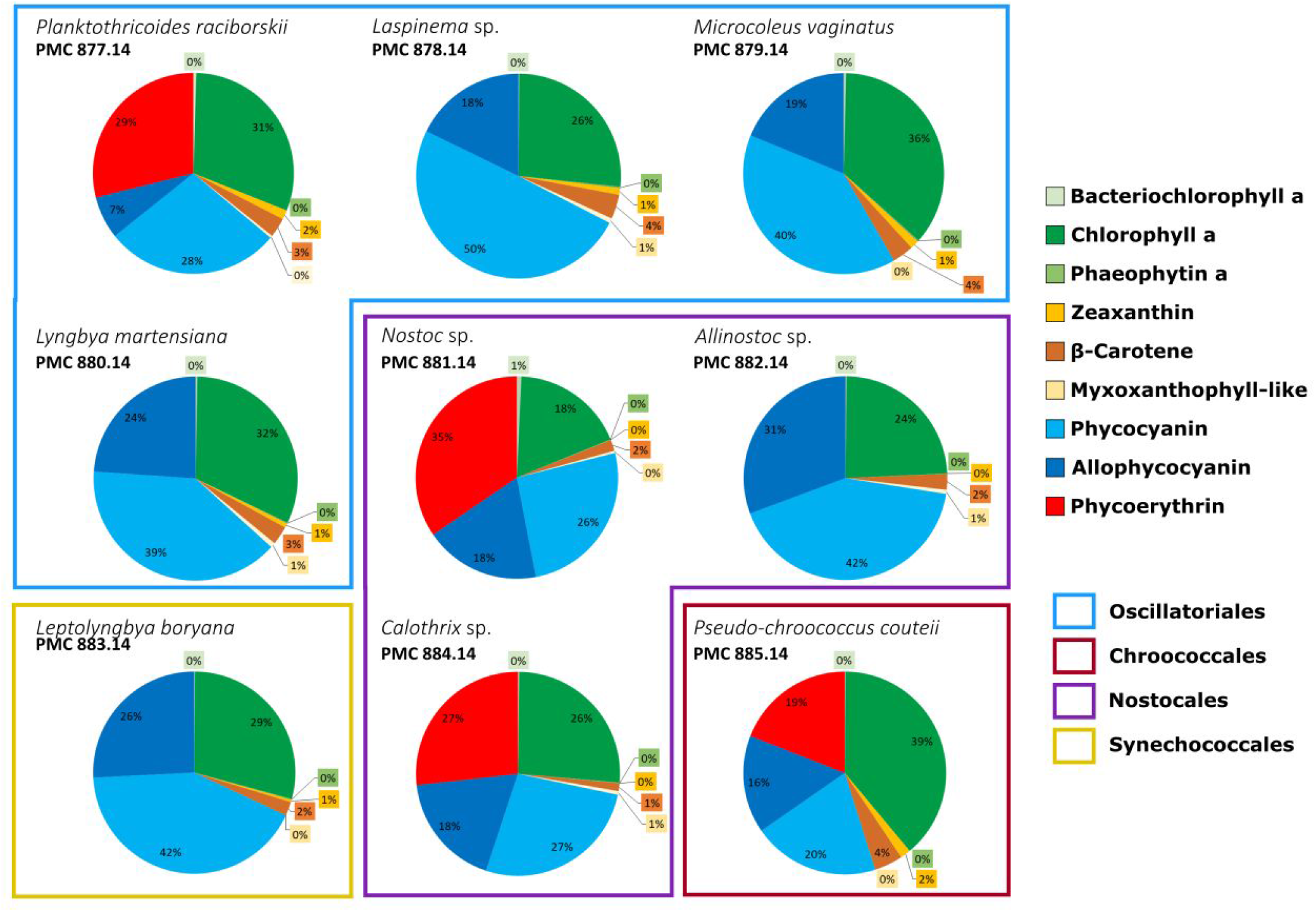
Pigments composition (%) of the nine studied strains, isolated from Thermes de Balaruc-Les-Bains. PMC: Paris Museum Collection, Museum National of Natural History.

**Figure S2.**
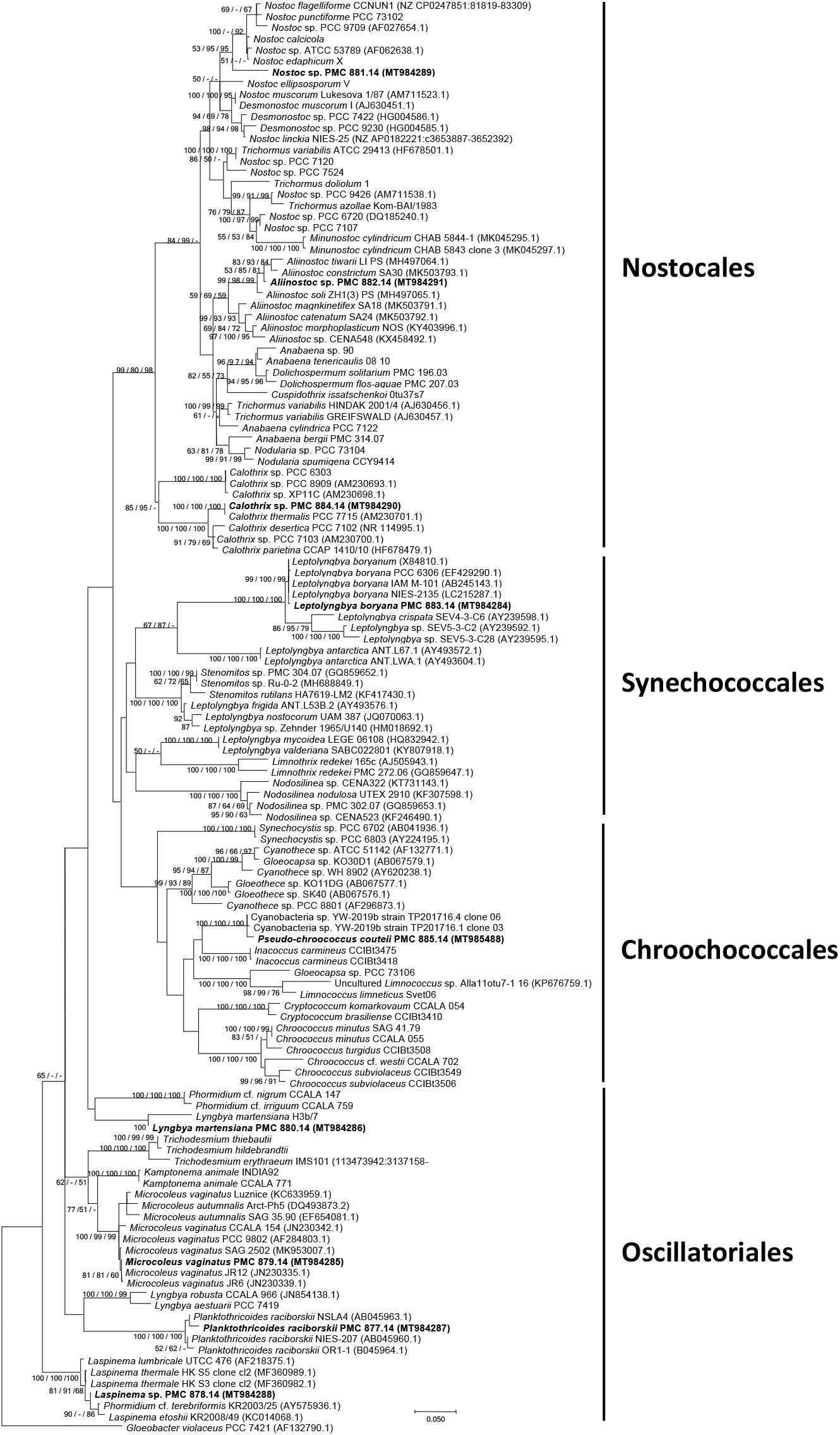
Consensus phylogenetic tree based on 16S rRNA gene sequences of representative cyanobacteria strains (129 sequences, 1446 aligned nucleotide positions, GTR+G+I model) belonging to the orders Oscillatoriales, Nostocales, Chroococcales, Synechococcales and the studied strains form Thermes de Balaruc-Les-Bains (in bold). *Gloeobacter violaceus* was used as outgroup. Numbers above branches indicate bootstrap support (>50%) from 1000 replicates. Bootstrap values are given in the following order: maximum likelihood / neighbor joining / maximum parsimony.

**Table S1.** List of GenBank accession numbers. PMC: Paris Museum Collection.

| Order | Species | Strain number | 16S | 16S -23S |
| --- | --- | --- | --- | --- |
| Chroococcales | *Pseudo-chroococcus couteii* | PMC 885.14 | - | MT985488 |
| Synechococcales | *Leptolyngbya boryana* | PMC 883.14 | MT984284 | - |
| Oscillatoriales | *Planktothricoides raciborskii* | PMC 877.14 | MT984287 | - |
|  | *Laspinema* sp. | PMC 878.14 | MT984288 | - |
|  | *Microcoleus vaginatus* | PMC 879.14 | MT984285 | - |
|  | *Lyngbya martensiana* | PMC 880.14 | MT984286 | - |
| Nostocales | *Nostoc* sp. | PMC 881.14 | MT984289 | - |
|  | *Aliinostoc* sp. | PMC 882.14 | MT984291 | - |
|  | *Calothrix* sp. | PMC 884.14 | MT984290 | - |

**Table S2.**
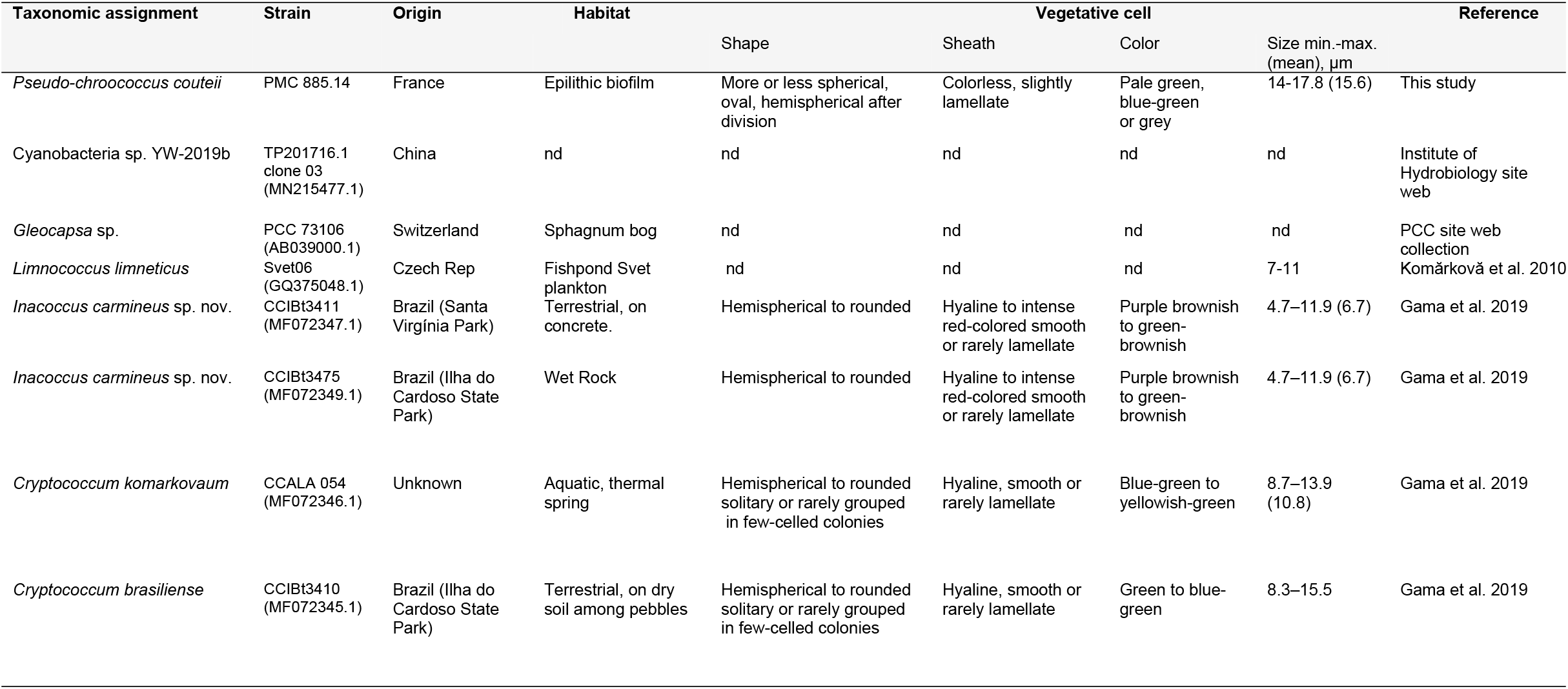
Comparison of the morphological features of *Pseudo-chroococcus couteii* PMC 885.14 strain with other Chroococcales (*Gleocapsa, Limnococcus, Inacoccus, Cryptococcum*) genera (Fig. 4). nd: not determined.

**Table S3.** Comparison of the morphological features of the *Leptolyngbya boryana* sp. PMC 883.14 strain with other *Leptolyngbya* species belonging to the same phylogenetical cluster (Fig. 8). nd: not determined.

| Taxonomic assignment | Strain | Origin | Habitat | Trichome |  |  |  |  | Vegetative cell |  |  | Reference |
| --- | --- | --- | --- | --- | --- | --- | --- | --- | --- | --- | --- | --- |
| | | | | Color | Sheath | Shape | Shape of terminal cell | Width min.-max. (mean) $\mu\text{m}$ | Shape | Width min.-max. (mean), $\mu\text{m}$ | Length min.-max. (mean), $\mu\text{m}$ | |
| *Leptolyngbya boryana* | PMC 883.14 | France | Freshwater, benthic, biofilms | Pale to bright blue-green | Thin, colourless, attached to the trichome | Straight, curved, distinctly constricted, | Rounded hemispherical | 1.5-2.4 | Moniliform, shorter than wide | 1.5-2.4 (2.1 ±0.24) length:width ratio = 0.75 | 0.9 - 2 (1.52 ±0.27) | This study |
| *Leptolyngbya boryana* | PCC 6306 (EF429290.1) | nd | Freshwater, metaphytic, lentic | Blue-green | Absent | Long, straight or wavy, distinctly constricted, false branching | Rounded | nd | More or less isodiametric | length:width ratio = 0.78 | length:width ratio = 0.78 | Chakraborty et al. 2019 |
| *Leptolyngbya boryana* | UAM 391 (JQ070062.1) | Spain Guadarrama river | Freshwater, benthic, biofilms | Intense blue-green or pale color | Hyaline, homogeneous sometimes thick, separated from the filament | Strongly constricted, false branching | Rounded | nd | nd | 2.1–3.2 | 1.2–3.6 | Loza et a. 2013 |
| *Leptolyngbya boryana* | IAM M-101 (AB245143.1) | Japan | Freshwater | nd | nd | nd | nd | nd | nd | nd | nd | nd |
| *Leptolyngbya boryana* | NIES-2135 (LC215287.1) | Japan | Freshwater | nd | nd | nd | nd | nd | nd | nd | nd | nd |
| *Leptolyngbya foveolarum* | 1964/112 (X84808.1) | nd | nd | nd | Presence of a sheath | Constrictions at the cross-walls | nd | nd | nd | 3.8 | 3.8 | Nelissen et al. 1996 |
| *Leptolyngbya boryanum* | PCC 73110 (X84810.1) | nd | Freshwater | nd | Variable | Constrictions at the cross-walls | nd | nd | nd | 1.9 – 2.6 | 1.9 – 2.9 | Nelissen et al. 1996 |

**Table S4.**
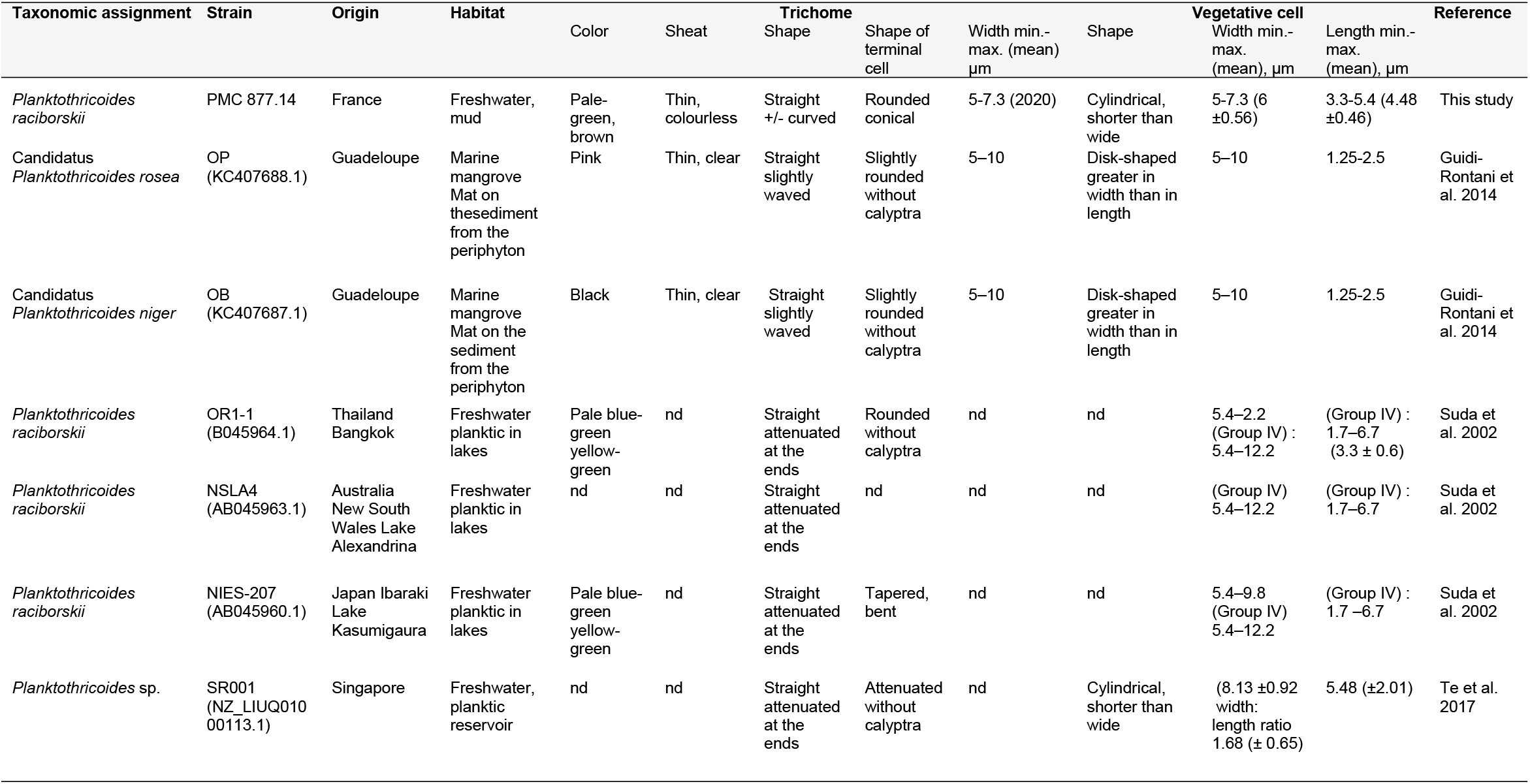
Comparison of the morphological features of the *Planktothricoides raciborskii* PMC 877.14 strain with other *Planktothricoides* species belonging to the same phylogenetical cluster (Fig. 9). nd: not determined.

**Table S5.** Comparison of the morphological features of the *Laspinema* sp. PMC 878.14 strain with other *Laspinema* and *Phormidium* species belonging to the same phylogenetical cluster (Fig. 9). nd: not determined.

| Taxonomic assignment | Strain | Origin | Habitat | Trichome |  |  |  |  | Vegetative cell |  |  | Reference |
| --- | --- | --- | --- | --- | --- | --- | --- | --- | --- | --- | --- | --- |
| | | | | Color | Sheath | Shape | Shape of terminal cell | Width min.-max. (mean) $\mu\text{m}$ | Shape | Width min.-max. (mean), $\mu\text{m}$ | Length min.-max. (mean), $\mu\text{m}$ | |
| *Laspinema* sp. | PMC 878.14 | France | Benthic on mud freshwater, brackish waters | Bright blue-green | Lacking sheath | Straight, nutated at the end | Obtuse-conical, rounded-conical, without calyptra | 3.8 - 4.8 | Shorter than wide | 3.8 - 4.8 (4.26 ±0.27) | 2.3 - 3.3 (2.87 ±0.28) | This study |
| *Laspinema thermale* | HK S5 clone 2 (MF360989.1) | Iran, Mahallat | Thermal spring | Blue-green or olive green | Thin, inconspicuous | Straight in natural environment forming benthic mats, slightly constricted on cross walls | Rounded, conical, shortly bent | 3 – 4 (5) | Shorter than wide | 3 – 4 (5) | 1 – 2,4 | Heidari et al. 2018 |
| *Laspinema lumbricale* | UTCC 476 (AF218375.1) | Antarctica | Saline pond on McMurdo Ice Shelf | nd | nd | nd | Rounded, straight | nd | nd | 5 – 6 (7) | 2 - 4 | Casamatta et al. 2005 |
| *Laspinema etoshii* | KR2008/49 (KC014068.1) | Namibia Etosha National Park, Okandeka Spring | Hyposaline water puddles springs and wet soil | Pale bright blue-green | Hyaline, colorless | Solitary, straight or curvy, not or slightly constricted | Round, conical; straight or bent without calyptra | nd | Mainly wider than long | 5.5 ±1.5 | 3.0 ±1.0 | Dadheech et al. 2013 |
| *Phormidium* cf. *okenii* | Led-Z (EU196644.1) | Czech Republ Lednice na Moravě park | a pond, benthos | Dark blue-green | Delicate colourless | Gradually narrowed towards end, distinctly constricted at cross-walls | Bent, elongated usually rounded | nd | Slightly shorter than wide or isodiametrical, sometimes longer than wide | 5.6-6.5 (3.7-7.5 ; 6.1) cell length/width ratio 0.5-0.8 | nd | Lokmer 2007 |
| *Phormidium* cf. *terebreformis* | KR2003/25 (AY575936.1) | Kenya, Hot spring, Lake Bogoria, | Hyposaline hot springs | nd | nd | Solitary, straight to undulating Constricted | Rounded without calyptra | nd | Mainly isodiametric | 5.5 ±0.5 | 3.0 ±1.0 | Dadheech et al. 2013 |

**Table S6.**
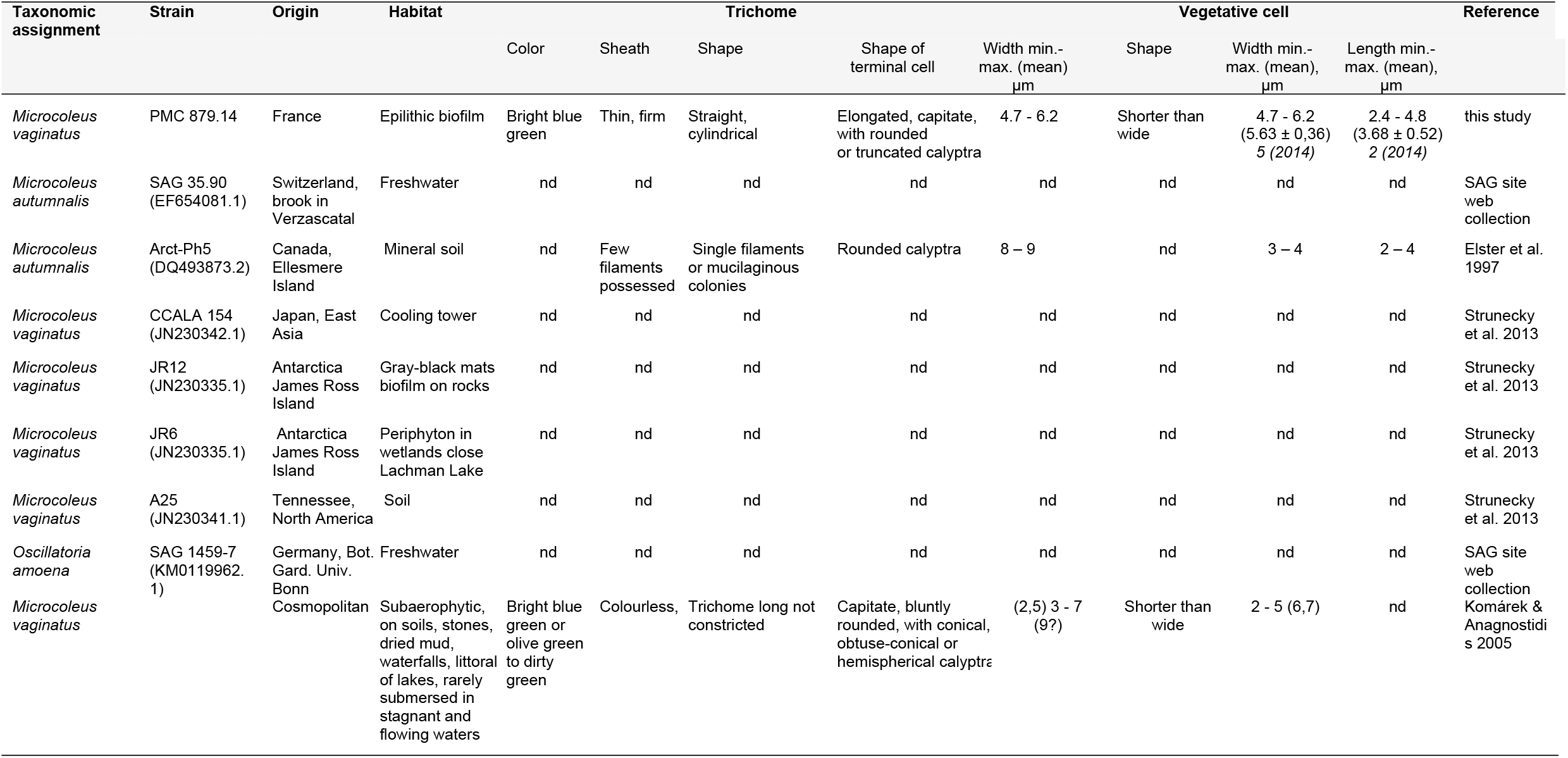
Comparison of the morphological features of the *Microcoleus vaginatus* PMC 879.14 strain with other *Microcoleus* species or strains belonging to the same phylogenetical cluster (Fig. 9). nd: not determined.

**Table S7.**
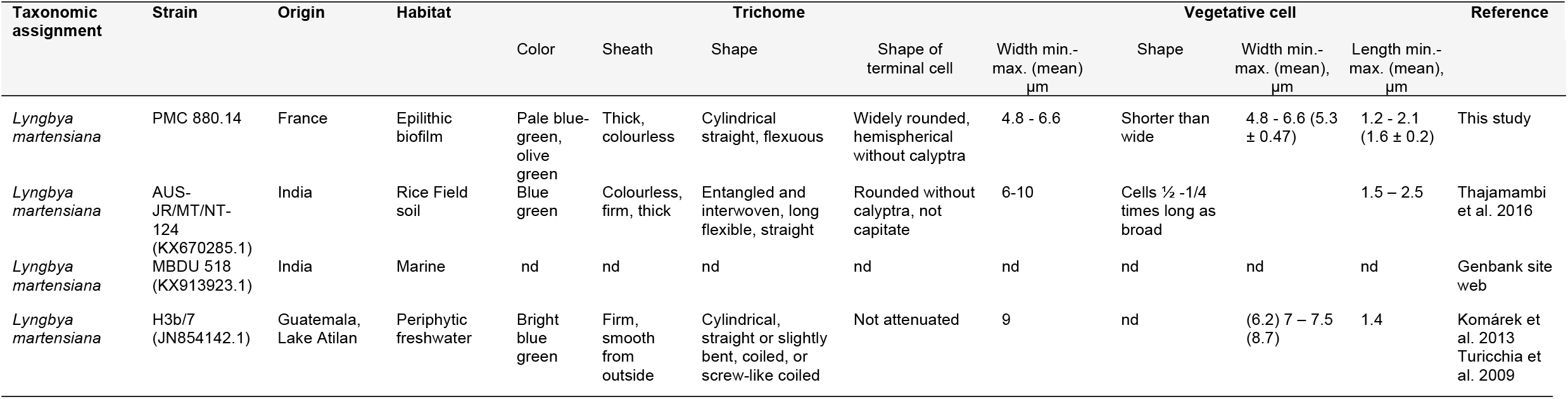
Comparison of the morphological features of the *Lyngbya martensiana* PMC 880.14 strain with other *Lyngbya martensiana* belonging to the same phylogenetical cluster (Fig. 9). nd: not determined.

**Table S8.**
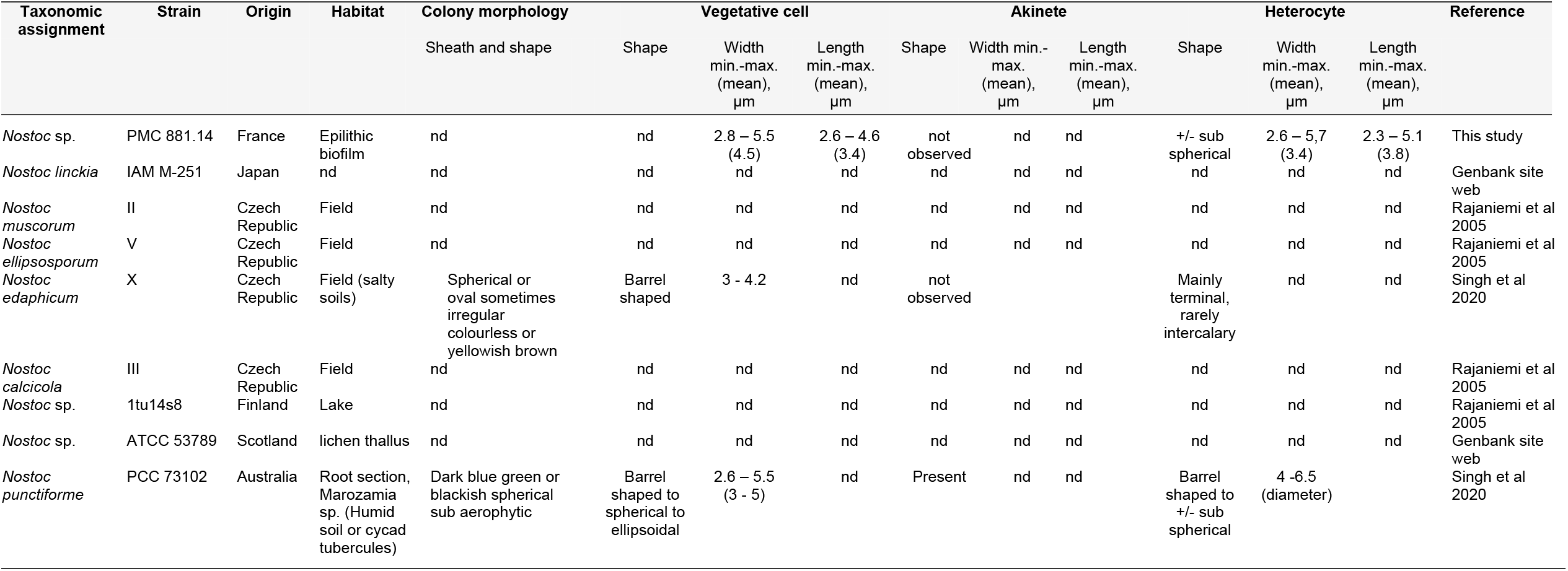
Comparison of the morphological features of the *Nostoc* sp. PMC 881.14 strain with other *Nostoc* species belonging to the same phylogenetical cluster (Fig. 14). nd: not determined.

**Table S9.**
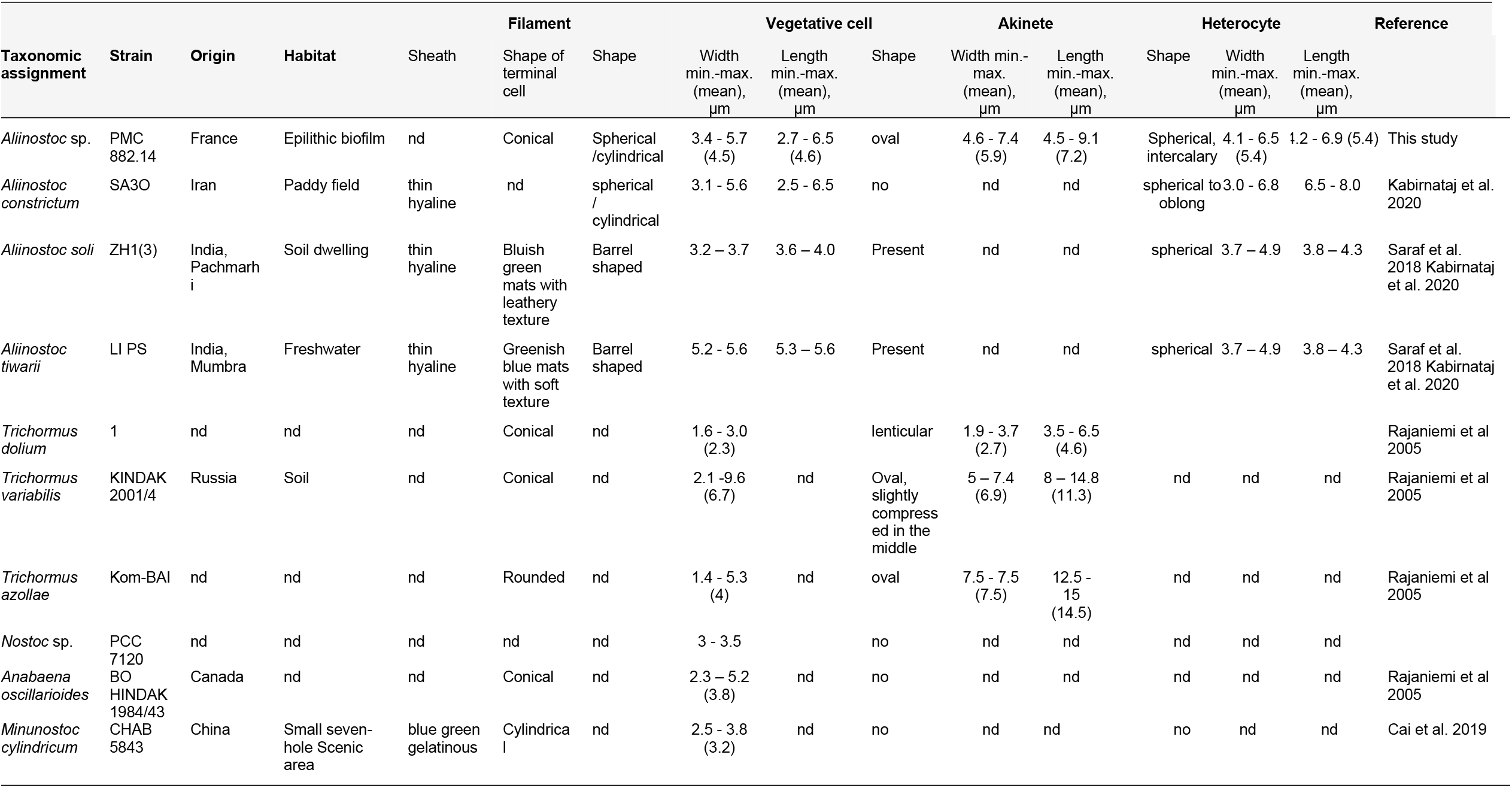
Comparison of the morphological features of the *Aliinostoc* sp. PMC 882.14 strain with other *Aliinostoc* and *Trichormus* species belonging to the same phylogenetical cluster (Fig. 14). no: not observed. nd: not determined.

**Table S10.** Comparison of the morphological features of the *Calothrix* sp. PMC 884.14 strain with other *Calothrix* species belonging to the same phylogenetical cluster (Fig. 14). nd: not determined.

| Taxonomic assignment | Strain | Origin | Habitat | Vegetative cell |  |  | Akinete |  | Heterocyte |  | Reference |
| --- | --- | --- | --- | --- | --- | --- | --- | --- | --- | --- | --- |
| | | | | Shape | Width min.-max. (mean), $\mu\text{m}$ | Length min.-max. (mean), $\mu\text{m}$ | Shape | Shape | Width min.-max. (mean), $\mu\text{m}$ | Length min.-max. (mean), $\mu\text{m}$ | |
| *Calothrix* sp. | PMC 884.14 | France | Epilithic biofilm | nd | 5.2 – 7.9 (6.6±0.7) | 4.8 – 6.2 (5.5±0.4) | not observed | hemispherical | 4.7-8.19 (6.2±0.9) | 3.48-7.33 (4.9±0.9) | This study |
| *Calothrix thermalis* | PCC 7715 | France | Thermal spring | Isodiametric to longer than wide cells | nd | nd | Present | nd | nd | nd | PCC site web collection |
| *Calothrix desertica* | PCC 7102 | Chile | Soil, fine desert sand | nd | nd | nd | nd | nd | nd | nd | PCC site web collection |
| *Calothrix* sp. | PCC 7103 | USA | Herbarium material of *Anacystis montana* | nd | nd | nd | nd | nd | nd | nd | PCC site web collection |
| *Calothrix parietina* | CCAP 1410/10 | England | Freshwater | nd | nd | nd | nd | nd | nd | nd | Genbank site web |

**Table S11.** Molecules selected for their antioxidant and/or anti-inflammatory properties (Demay et al. 2019) related to the genera of cyanobacteria isolated from Thermes de Balaruc-Les-Bains. MAAs: Mycosporine-like amino acids.

| Targeted molecule families | Characteristics | Bioactivities | Genus producers | References |
| --- | --- | --- | --- | --- |
| **Aeruginosins** | Peptide<br>Linear<br>NRPS<br>Hydrosoluble | No cytotoxicity<br>Anti-inflammatory activity<br>Protease inhibitor (trypsin, thrombin, plasmin) | *Nostoc* sp. Lukešová 30/93 | [1–4] |
| **Carotenoids** | Terpenoid<br>Pigment<br>Liposoluble | Antioxidant<br>Sunscreen | All cyanobacteria | [5–10] |
| **Chlorophylls** | Substituted tetra-pyrrole porphyrin molecule<br>Pigment<br>Liposoluble | Antioxidant<br>Antimutagenic<br>Chemopreventive<br>Photosensitizing act.<br>Cancer preventive agent | All cyanobacteria | [11–13] |
| **Honaucins** | Lactone<br>Linear<br>Liposoluble | Anti-inflammatory activity<br>No antioxydant act.<br>Quorum sensing inhibition | *Leptolyngbya crossbyana* HI09-1 | [14,15] |
| **Mycosporine-like amino acids (MAAs)** | Cyclohexenone-amino acid<br>Hydrosoluble | Antioxidant<br>UV-absorbing<br>UV-protective act.<br>Sunscreen<br>Antiphotaging | *Calothrix parietina*<br>*Calothrix* sp.<br>*Leptolyngbya* sp.<br>*Lyngbya aestuarii*<br>*Lyngbya* sp. CU2555<br>*Lyngbya* sp.<br>*Microcoleus chthonoplastes*<br>*Microcoleus paludosus*<br>*Microcoleus* sp.<br>*Nostoc commune*<br>*Nostoc microscopicum*<br>*Nostoc punctiforme* ATCC 29133<br>*Nostoc* sp. | [5,16–25] |
| **Phycocyanins** | Protein<br>Pigment<br>Hydrosoluble | Antioxidant<br>Anti-inflammatory<br>Neuroprotective effects<br>Hepatoprotective effects | All cyanobacteria | [26–29] |
| **Phycoerythrins** | Protein<br>Pigment<br>Hydrosoluble | Antioxidant<br>Moderate aging | Several strains belonging to all orders | [30–32] |
| **Scytonemins** | Alkaloid<br>Pigment<br>Liposoluble | Anti-inflammatory<br>UV protection<br>Inhibit cell cycle kinases<br>Antiproliferative<br>Induction of autophagic death | *Calothrix crustacea*<br>*Calothrix parietina*<br>*Calothrix* sp.<br>*Chroococcus* sp.<br>*Lyngbya aestuarii*<br>*Lyngbya* sp.<br>*Lyngbya* sp. CU2555<br>*Nostoc commune* Vauch<br>*Nostoc microscopicum*<br>*Nostoc parmelioides*<br>*Nostoc pruniforme*<br>*Nostoc punctiforme* PCC 73102, ATCC 29133 | [16,33–38] |

